# Optimised fluorescence-activated nuclei sorting for epigenomic analysis of cortical cell types

**DOI:** 10.64898/2025.12.22.695789

**Authors:** Barry Chioza, Stefania Policicchio, Joe Burrage, Georgina E T Blake, Rosemary A. Bamford, Alice Franklin, Darren Soanes, Philippa M. Wells, Ann Babtie, Marina Flores Payan, Jonathan P. Davies, Anthony Klokkaris, Emma M Walker, Joy N. Ismail, Paulina Urbanaviciute, Sarah J. Marzi, Eilis Hannon, Jonathan Mill, Emma L Dempster

**Author notes:** These authors contributed equally to this work (Barry Chioza, Stefania Policicchio). These authors co-supervised this work (Jonathan Mill, Emma L Dempster).

## Abstract

Increased understanding of the functional complexity of the genome has led to growing recognition of the role of non-sequence-based regulatory variation in disorders of the human central nervous system. Most genomic analyses of the brain are limited by the use of bulk tissue, which comprises a heterogeneous mix of different neural cell types with distinct epigenetic profiles, thereby limiting the ability to attribute regulatory changes to specific cell populations. Given the limited availability of human post-mortem tissue resources and the importance of integrating multi-omic data from the same samples, there is a critical need for methods that enable parallel, cell-type–resolved genomic profiling. We present optimised protocols using fluorescence-activated nuclei sorting (FANS) to isolate nuclei from different human and mouse brain cell types for downstream multi-omic analysis. Our approach enables the robust purification of neuronal, oligodendrocyte, microglial and other glial-origin nuclei from both adult and fetal brain tissue. We demonstrate that FANS-isolated nuclei are compatible with a wide range of genomic assays, including profiling of DNA modifications, histone modifications, chromatin accessibility, and gene expression. This protocol maximises the utility of limited post-mortem tissue resources and provides a unified workflow for comprehensive, cell-type–specific interrogation of molecular mechanisms involved in the brain.

## Introduction

Despite success in identifying genetic risk factors for neuropsychiatric (1), neurodegenerative disease (2) and other central nervous system conditions, little is known about the functional mechanisms involved in pathogenesis. Increased understanding of the functional complexity of the genome has led to the recognition of the role of non-sequence-based regulatory variation in health and disease. Of particular interest are epigenomic processes which act to developmentally control gene expression via modifications to DNA, histone proteins and chromatin. These mechanisms play a critical role in determining cell-type-specific patterns of gene transcription in the human brain, and epigenetic variation has been associated with conditions including Alzheimer’s disease (AD) (2), schizophrenia (3) and autism (4). Analyses of transcriptional and epigenomic variation in human post-mortem brain tissue are often performed on ‘bulk’ tissue, which comprises a complex mix of different neural cell types. Because cellular heterogeneity is critical to consider in studies of disorders that affect specific cell populations – for example, neurodegenerative diseases such as AD are characterised by extensive neuronal loss in conjunction with glial cell activation and proliferation (5) - the development of methods to quantify genomic variation in specific cell-types is important.

Recent advances in single cell sequencing technologies have enabled researchers to quantify transcriptomic variation in individual neural cells (6, 7), but their application remains constrained by cost, scale, and the difficulty of profiling multiple genomic modalities from the same sample. Recently, methods using fluorescence-activated nuclei sorting (FANS) have been developed to purify nuclei populations enriched for neurons using an antibody to NeuN, a robust marker of post-mitotic neurons (8), prior to the quantification of DNA methylation, histone modifications and chromatin accessibility (9, 10). This method has been extended to the interrogation of other neural cell types, including oligodendrocytes (11) and microglia (12). However, existing protocols typically isolate nuclei from only one or two cell types at a time and are not optimised for parallel multi-omic profiling across multiple neural populations isolated from small amounts post-mortem tissue.

To address these challenges, we have developed and validated a set of FANS protocols for the quantification of multiple markers of gene regulation (including DNA modifications, chromatin accessibility, H3K27 acetylation and gene expression) across multiple neural cell populations. Our primary protocol involves the simultaneous isolation of nuclei populations enriched for i) neurons (using an antibody to NeuN, a robust marker of post-mitotic neurons (8)), ii) oligodendrocytes (using an antibody to SOX10, a transcription factor involved in the differentiation of oligodendrocytes (13)) and iii) microglia (using an antibody to interferon regulatory factor 8 (IRF8), a transcription factor that plays critical roles in the regulation of lineage commitment and is upregulated in activated microglia (14)) from a relatively small quantity (∼300mg) of cryopreserved human post-mortem cortex tissue. Additional protocols are provided for the purification of different neuronal subtypes (e.g. excitatory (SATB2^+^) neurons) and for the isolation of purified nuclei from mouse cortex.

We validate the identity and purity of these sorted populations using three different transcriptomic assays (qPCR of cell marker genes, RNA-seq of bulk purified nuclei populations and single nucleus RNA-seq (snRNA-seq)). We further demonstrate that nuclei isolated using our protocols are compatible with a wide range of downstream genomic assays the quantification of DNA methylation (5mC) using the Illumina EPIC array, 5mC and hydroxymethylcytosine (5hmC) using Oxford Nanopore Technologies long-read sequencing (ONT), chromatin accessibility using the Assay for Transposase-Accessible Chromatin in combination with highly-parallel sequencing (ATAC-seq) and the histone modification H3K27ac using CUT&Tag sequencing. Applying these methods to profile frozen tissue samples from clinical donors and rodent models maximises the utility of these limited tissue resources. Our protocols, which are available as a resource on protocols.io, represent versatile methods for the dissection of cell-type-specific regulatory mechanisms in the brain, providing new opportunities to investigate the molecular pathways underlying disorders of the central nervous system.

## Material and Methods

### Fluorescence-activated nuclei sorting (FANS) from post-mortem cortex

We present optimised FANS protocols for the immunolabelling and purification of nuclei from post-mortem human and mouse cortex. This workflow supports the parallel isolation of distinct nuclear populations, including neuronal-enriched (NeuN^+^ or SATB2^+^), oligodendrocyte-enriched (SOX10^+^), and microglia-enriched (IRF8^+^) nuclei from the same sample, enabling downstream multi-omic profiling. Detailed versions of all FANS protocols can be found in Supplementary Files Protocol_S1-3 and are also available on protocols.io via the following links:

- Isolating neuron-enriched, oligodendrocyte-enriched, microglia-enriched and astrocyte-enriched nuclei populations from human postnatal cortex: https://dx.doi.org/10.17504/protocols.io.36wgq4965vk5/v2
- Isolating excitatory neuron-enriched nuclei populations from human fetal and postnatal cortex: https://dx.doi.org/10.17504/protocols.io.n92ldz9d8v5b/v1
- Isolating neuron-enriched and glia-enriched nuclei populations from mouse cortex: https://dx.doi.org/10.17504/protocols.io.dm6gpbwndlzp/v2

### Summary of key steps

Figure 1 provides a graphical overview of our FANS workflow. All samples used in this study are drawn from ongoing research projects using flash-frozen prefrontal cortex tissue obtained from sources. Human prefrontal cortex (PFC) tissue was obtained from several registered brain banks. Subjects provided written consent for brain donation during life, and all tissue collections were conducted and stored in accordance with legal and ethical guidelines. Further details are provided in the acknowledgements. Mouse frontal cortex tissue was dissected from 30 adult (1.5–12 months) wildtype C57BL/6J mice as described in Clifton *et al* (2025) (15). Approximately 300 mg of frozen postnatal human cortex, 200 mg of frozen fetal human cortex, or 100 mg of frozen mouse cortex tissue was homogenised in 2 mL of lysis buffer using a pre-chilled Dounce homogeniser. Nuclei were isolated via ultracentrifugation over a sucrose cushion. For flow cytometry, 150 µL of the nuclei suspension was reserved as an unstained control, transferred into a new 2 mL tube, mixed with 2 μL of Hoechst 33342 (Abcam, Cat# ab228551), and the volume adjusted to 1 mL with sucrose buffer (SB). The remaining nuclei were immunostained with antibody panels, depending on the species (human vs. mouse), developmental stage (fetal vs. postnatal), and target cell type (see **Table 1** for optimised FANS-compatible antibodies). Our main panels consist of the following antibody combinations:

‒ Adult Human: (NeuN–Alexa Fluor 488; SOX10–NL577; IRF8–APC)
‒ Fetal Human: (SATB2–Alexa Fluor 488 (replacing NeuN))
‒ Mouse:(NeuN–Alexa Fluor 488; PU.1-PE)

**Figure 1.**
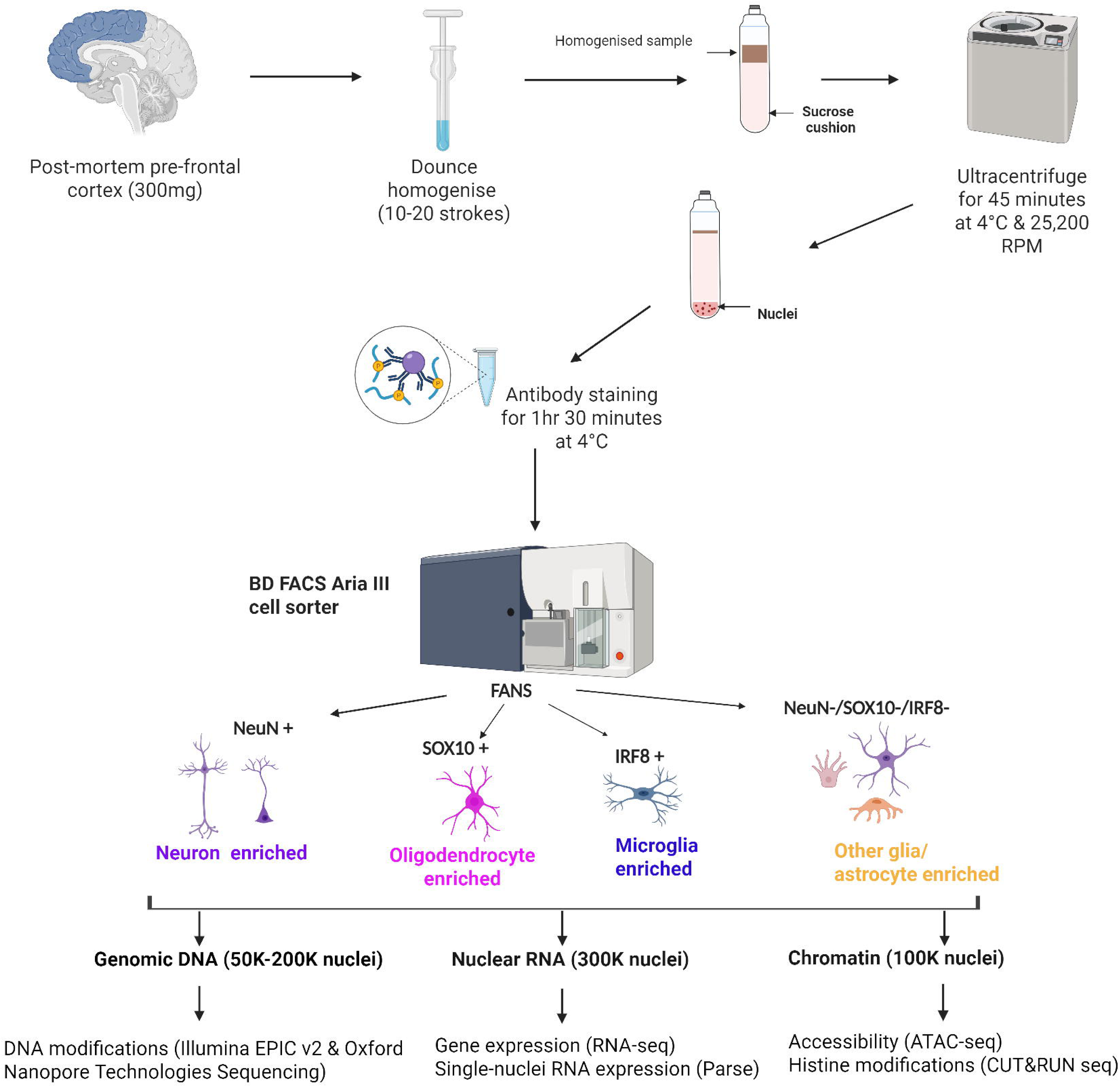
Schematic showing an overview of the process for generating multi-omic data from purified nuclei populations from tissue to different data types. Figure created with BioRender.com.

**Table 1.**
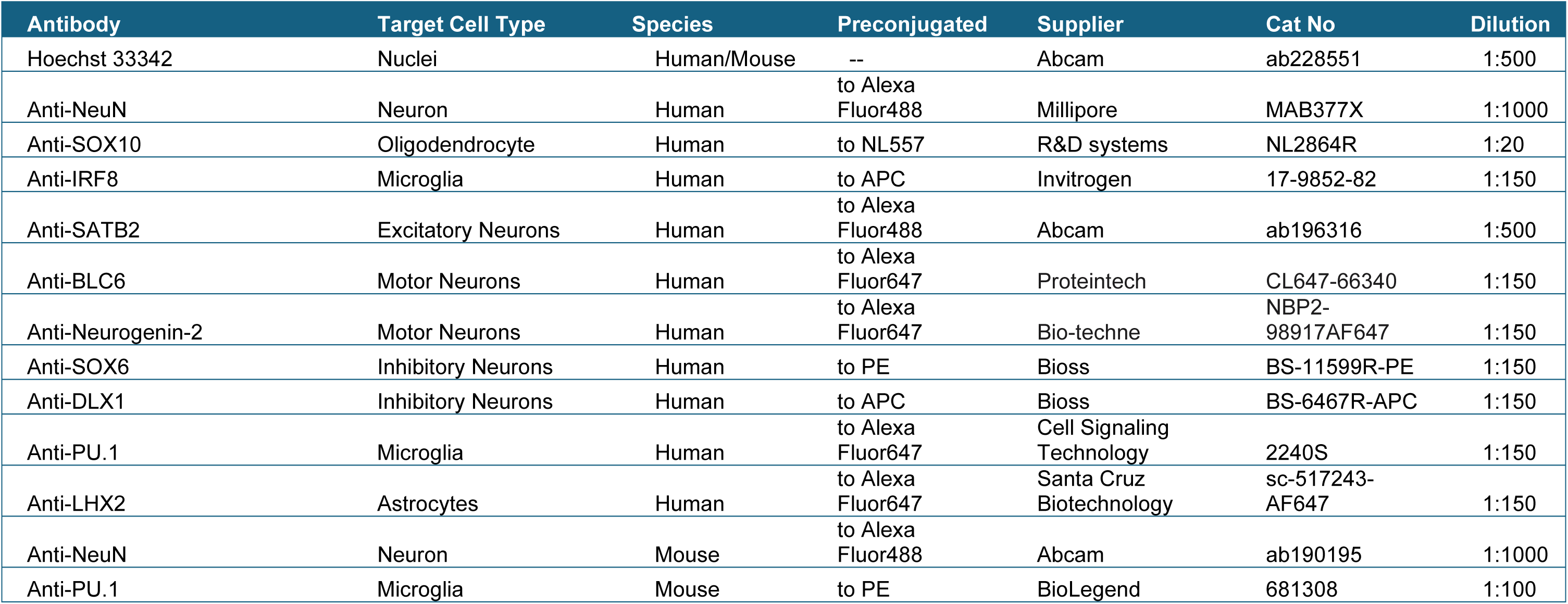
Details of optimised FANS-compatible antibodies.

Samples were incubated for 1.5 hours on a rotating platform at 4 °C in the dark. After incubation, samples were centrifuged (1000 × g, 5 min, 4 °C), the supernatant discarded, and nuclei pellets resuspended in SB (500 µL for unstained, 1–1.5 mL for stained, depending on pellet density) using wide-bore tips. Samples were kept on ice throughout sorting using the BD FACSAria III. A 100 μm nozzle was used, and the event rate was maintained at ≤ 3000 events/sec. Gating strategies were optimised to remove debris and identify target populations (Figure 2a-e**, Figure S1 and Figure S2**). Sorted nuclei were collected for downstream multi-omic analysis, including RNA-seq, DNA methylation profiling, ATAC-seq, and CUT&Tag. Minor adaptations were required for mouse tissue, including reduced ultracentrifugation time (30 min), substitution of PU.1 for IRF8 for microglial labelling, and use of alternative NeuN and SOX10 antibodies due to cross-reactivity. Representative gating strategies for mouse cortex are shown in **Figure S3**.

**Figure 2.**
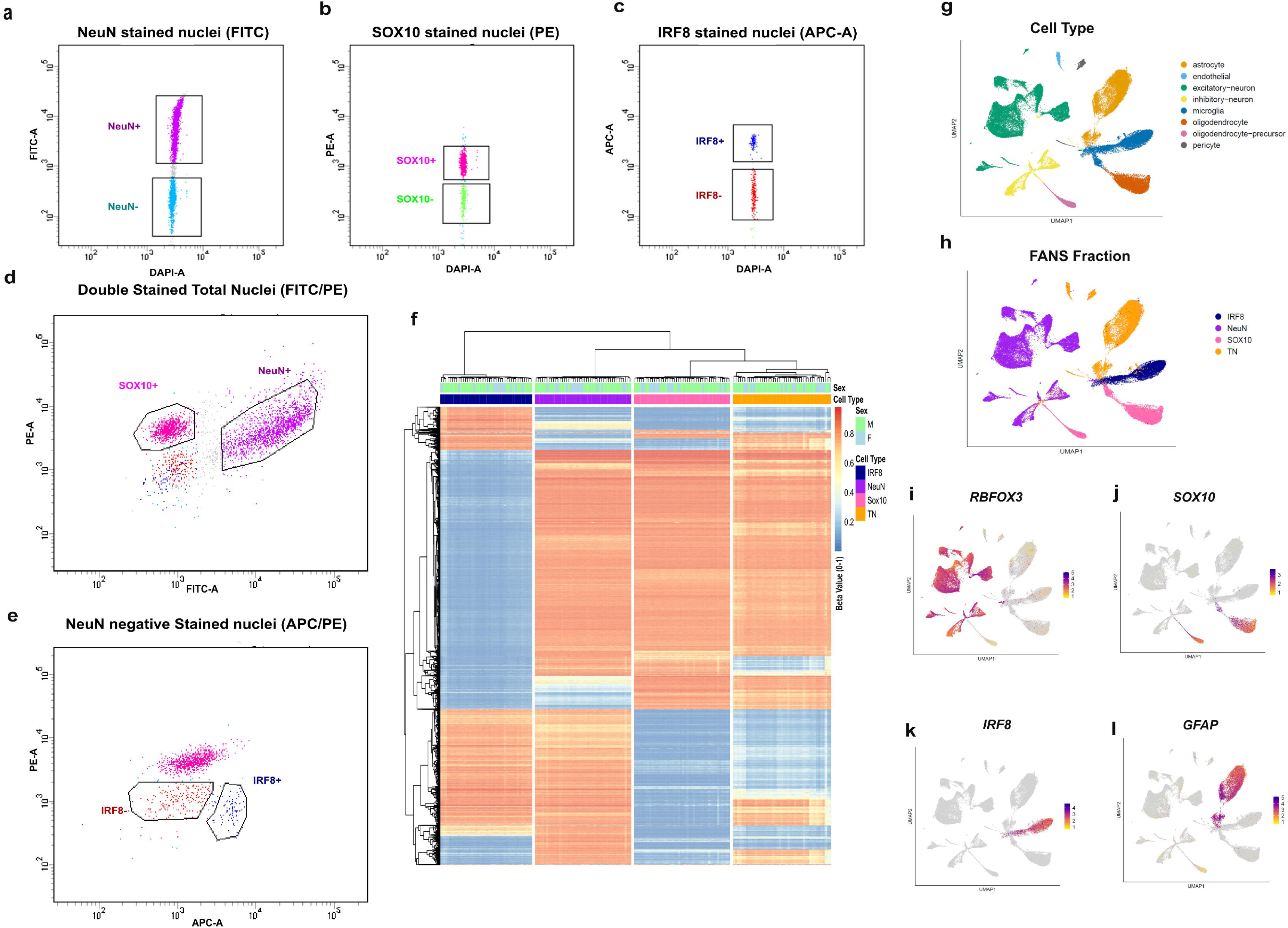
FANS gating strategy for isolating discrete nuclei populations and validating cell-type specificity using multi-omic profiling. (a–e) Sequential gating was used to isolate major neural nuclei populations based on antibody fluorescence intensity. Scatterplots show nuclei positive for (a) NeuN–Alexa Fluor 488 (purple; neurons), (b) SOX10–NL577 (dark pink; oligodendrocytes), and (c) IRF8–APC (dark blue; microglia). (d–e) Combined gating of all three markers enabled classification of nuclei into four mutually exclusive groups: NeuNL neurons, SOX10L oligodendrocytes, IRF8L microglia, and triple-negative (NeuNL/SOX10L/IRF8L) nuclei, which are enriched for astrocytes. (f) DNA methylation profiling of FANS-sorted nuclei demonstrates strong cell-type-specific epigenetic signatures. Heatmap and hierarchical clustering are based on the top 1,000 most variable CpG sites measured using the Illumina EPIC v2 array. After quality control, 889,069 CpG sites were retained across 47 individuals (14 female, 33 male; ages 6–91 years), with four nuclei fractions profiled per individual (NeuN: n=44; SOX10: n=44; IRF8: n=42; triple-negative: n=45). (g–k) Single-nucleus RNA-seq (snRNA-seq) of approximately 100,000 nuclei per donor, processed using the Parse Biosciences Evercode low-input fixation workflow, further confirms the specificity of FANS-isolated populations. Four sorted fractions were analysed per donor: NeuN⁺ (neuronal), SOX10⁺ (oligodendrocyte), IRF8⁺ (microglial), and triple-negative (residual nuclei negative for all three markers). (g) UMAP of predicted cell identities inferred using the Seurat workflow (h) UMAP clusters correspond to the FANS-defined populations. (i–l) Expression of canonical marker genes validates each cluster: (i) *RBFOX3* in NeuNL nuclei, (j) *SOX10* in SOX10L nuclei, k) *IRF8* in IRF8L nuclei, and (l) *GFAP* in the triple-negative, astrocyte-enriched population.

### RNA isolation and qPCR analysis

When collecting nuclei for RNA processing, a slight modification was made to the FANS protocol to maximise RNA integrity. Ribolock RNase-inhibitor (Thermo Scientific, EO0382) was added to the lysis buffer (0.2 U/ml) and staining buffer (0.2 U/ml). 300,000 nuclei were sorted into 1 mL of pre-chilled TRIzol™ LS Reagent (Invitrogen™, Cat N#11588616) and snap frozen on dry ice. RNA was extracted from the separated nuclei suspension using the Direct-zol™ RNA MicroPrep column kit (Zymo Research, Cat #R2060). Complementary DNA (cDNA) was reverse transcribed using the Invitrogen VILO cDNA synthesis kit (Life Technologies, Cat # 11754050) in 20 μL reactions according to the manufacturer’s instructions. Quantitative RT-PCR was performed in duplicate using the QuantStudio 12 K Flex (Applied Biosystems) in conjunction with the TaqMan low-density array (TLDA) platform using off-the-shelf pre-optimised assays (**Table S1**). Marker genes included *RBFOX3* (neurons), *OLIG2* (oligodendrocytes), *GFAP* (astrocytes), *CD68* (microglia) and five housekeeping genes (*ACTB*, *EIF4A2*, *GAPDH*, *SF3A1*, *UBC*) identified as being most stably expressed in the brain using GeNORM (Primer Design, Southampton, UK). PCR cycling conditions were 50 °C for 2 min, 94.5 °C for 10 min, and 45 cycles of 97 °C for 15 s and 60 °C for 1 min. Raw Ct data underwent stringent QC, removing reactions with high replicate variability (Ct difference > 0.5). Gene abundance was calculated using the comparative Ct (ΔΔCt) method, normalised to the geometric mean of five housekeeping genes (*ACTB*, *EIF4A2*, *GAPDH*, *SF3A1*, *UBC*). Normalised values were log2-transformed. Expression levels for each sorted population were expressed as fold-change relative to unsorted (total) nuclei from the same donor.

### Bulk RNA-seq of nuclei populations

RNA-seq libraries were created from 10 ng RNA input using the NEBNext® Ultra™ II RNA Library Prep Kit for Illumina (New England Biolabs). Libraries were pooled and sequenced on an Illumina NovaSeq 6000 (150 bp paired-end, ∼80 million reads/sample). Following sequencing, adaptor sequences and low-quality bases (mean quality < 22) were trimmed using fastp v0.23.1 (16); reads < 75 bp post-trimming were discarded. Trimmed reads were aligned to the GRCh38 primary assembly using STAR v2.7.10b (17) with GENCODE Release 44 annotations. Gene-level counts were obtained using STAR’s *GeneCounts* mode. Downstream analyses were performed using DESeq2 v1.36.0 (18) to identify differential transcript abundance between sorted populations.

### Single-nucleus RNA sequencing (snRNA-Seq) using Parse Biosciences

100,000 sorted nuclei representing each nuclei fraction (NeuN^+^, SOX10^+^, IRF8^+^, and triple-negative) were fixed with the Parse Evercode Low-Input Nuclei Fixation Kit (v3). Approximately 6,250 nuclei per sample were targeted for recovery. Libraries were prepared with the Evercode WT Kit (v3) and assessed using the Agilent D5000 High Sensitivity ScreenTape assay. Sequencing libraries were subsequently pooled and run on a NovaSeq 6000 system (100 bp, paired-end reads, at a sequencing depth of 20,000 reads per nuclei). Trailmaker™ (v1.5.1) (Parse Biosciences) was used to align reads to GRCh38 and generate filtered count matrices (these exclude barcodes with fewer than 10 transcripts and those that do not meet the barcode rank plot threshold). Potential doublets were identified and excluded using the *scDblFinder* algorithm. Subsequent data analysis and visualisation used the Seurat R package (v5.2.1). Nuclei with at least 200 transcript counts and less than 10% mitochondrial transcripts, and genes detected in at least 5 nuclei were retained. Data from each sample were log-normalised, scaled, and the top 4000 variable features selected before dimensionality reduction using principal component analysis (PCA). Samples were integrated using Harmony to correct for batch effects, and clusters were identified using the Leiden community-detection algorithm. A Uniform Manifold Approximation and Projection (UMAP) embedding was generated for visualisation and interpretation of the resulting cellular structure. Clusters were manually annotated as major neural cell types using marker gene expression patterns(19–21) after excluding nuclei with fewer than 400 detected genes in a second round of filtering.

### Single-nucleus RNA sequencing (snRNA-Seq) using 10X Genomics

Sorted nuclei representing two nuclei fractions (NeuN^+^, SOX10 and total^)^ were inspected for debris and quantified with a haemocytometer before single-nucleus capture using the 10x Genomics Single-Cell 3′ platform, targeting ∼3,000 nuclei per sample. Library preparation was carried out following the 10x Chromium Next GEM Single Cell 3′ protocol (v3.1). Quantification, quality assessment, and fragment size analysis of cDNA and final libraries were performed using the D5000 High Sensitivity ScreenTape assay and reagents (Agilent Technologies). Sequencing libraries were subsequently pooled and run on a NovaSeq 6000 system. Raw sequencing data were processed using the Cell Ranger pipeline (v3.1.0). This included demultiplexing, generation of FASTQ files, alignment of reads to a pre-mRNA GRCh38 reference, and quantification of unique molecular identifiers (UMIs). Downstream data processing and visualisation were performed in Seurat R package (v3.6) (22). Nuclei with at least 200 transcript counts and less than 5% mitochondrial transcripts were retained. Data from each sample were log normalised and scaled, before dimensionality reduction using principal component analysis (PCA), cluster identification and tSNE embeddings generation (22). Clusters were manually annotated using marker gene expression patterns (19–21). These analyses were used to confirm successful enrichment of neuronal, oligodendrocyte, microglial, and triple-negative populations isolated by FANS.

### Illumina DNA methylation EPIC array

Approximately 50,000 nuclei per sample were processed using the Zymo EZ-96 DNA Methylation-Direct Kit. Bisulfite-converted DNA was hybridised to the Illumina Infinium MethylationEPIC 850K array (human) or MouseMethylation (mouse) array following manufacturer instructions. Samples from each donor (four fractions) were randomised across chips to minimise batch effects. Arrays were scanned on an Illumina HiScan system (Illumina, CA, USA). Raw IDAT files were processed using the bigmelon package in R (23), following the quality control pipeline described in detail by Walker et al. (2024) (24). For human samples, cell-type purity was further evaluated using a brain-specific cell deconvolution algorithm as previously described (25). Following quality control, data were normalised with the *dasen* method (26), applied separately for each cell type to prevent cross-population bias. Detailed scripts for these analyses are available at https://github.com/ejh243/BrainFANS.

### Oxford Nanopore Sequencing (ONT)

DNA was extracted from ∼500,000 nuclei were treated with RNase A and Proteinase K. DNA was extracted with a bead solution (PEG 8000 20%, 2.5M NaCl) and purified using 0.1x Ampure XP beads. DNA was eluted in nuclease-free water.

DNA quantity was assessed using a Qubit fluorometer, and size distribution/integrity was evaluated using Agilent Genomic DNA screentape. DNA was sheared to 5 kb using a Covaris g-TUBE, libraries were prepared according to ONT protocols and sequenced on an ONT PromethION using R10.4.1 flow cells. Oxford Nanopore sequencing data were processed using the workflow available at https://github.com/dmsoanes/ont-dna-modification/. Briefly, basecalling, including 5mC and 5hmC detection from POD5 files, was performed using Dorado v0.6.0 with the “dna_r10.4.1_e8.2_400bps_sup@v4.3.0” model (27). Reads were filtered at a minimum Q-score threshold of 10 and aligned to the GRCh38 reference genome (GENCODE Release 44).

### Chromatin accessibility profiling (ATAC-seq)

Approximately 100,000 nuclei were pelleted (1,000 × g, 5 min, 4 °C), resuspended in BAMBANKER™ Cryopreservative, and frozen at −20 °C. For library preparation, nuclei were thawed, washed with PBS, and processed following the Omni-ATAC protocol (28), including transposition and PCR amplification using NEBNext 2X Master Mix and custom Nextera primers. Libraries were purified using double-sided SPRI bead selection, quantified using the NEBNext Library Quant Kit, pooled equimolarly, and sequenced on an Illumina NovaSeq 6000/X (50 bp paired-end, ∼50 million reads/sample). ATAC-seq data were processed using a bespoke pipeline (https://github.com/ejh243/BrainFANS). FastQC (29) was used for initial quality control, and adapter or low-quality bases were removed using fastp. Reads were aligned to the GRCh38 reference genome using Bowtie2 (30), and low-quality, duplicate, unpaired, or mitochondrial reads were removed using SAMtools (31) and Picard (32). ATAC-seq–specific quality metrics, including nucleosome patterning, nucleosome-free region rates, alignment rates, and library complexity, were quantified using phantompeakqualtools and Picard following ENCODE ATAC-seq standards. Sample identity was confirmed using sex-chromosome–based sex prediction and VerifyBamID comparisons against donor genotype data(33). Peaks were identified for individual samples using MACS3 (34) with BAMPE mode,--nomodel flag and an FDR threshold of 0.05. Consensus peak sets for each cell type were identified by calling peaks on a merged BAM file comprising all QC-passing samples using MACS3 with a false discovery rate (FDR) threshold of 0.001. These merged peak sets were used as a reference and retained only if they were also detected in at least one individual sample. Read counts across consensus peak sets were quantified using Rsubread (35) and normalised to account for sequencing depth, library size, and technical variation.

### H3K27ac profiling (CUT&Tag)

For each assay, 100,000 nuclei were collected, pelleted (1,000 × g, 5 min, 4 °C), resuspended in BAMBANKER™, and stored at −20 °C. Nanobody-based CUT&Tag was performed according to published protocols (36, 37). Libraries were quantified using the NEBNext Library Quant Kit, pooled equimolarly, and sequenced on an Illumina NovaSeq 6000/X (100 bp paired-end, ∼20 million reads/sample). CUT&Tag sequencing data were processed using the nf-core/cutandrun pipeline (v3.2.2) (38). Adapter trimming was performed with TrimGalore (39), and reads were aligned to the GRCh38 reference genome using Bowtie2 (30). PCR duplicates were removed with Picard (32), and linear amplification duplicates were removed using a custom script. Sample identity was validated using sex chromosome read proportions and VerifyBamID (33) using donor genotype data. Peaks were called for merged cell-type–specific BAM files using MACS2 v2.2.9.1 (34) in narrow peak mode with the parameters --slocal 1000 --llocal 30000, --nomodel, and a q-value threshold of 1×10⁻L. Peaks overlapping ENCODE blacklist regions or non-canonical chromosomes were removed prior to downstream analysis. Read quantification was performed using featureCounts v2.0.1 (40). Cell-type purity for each remaining sample was assessed using CHAS v0.99.9 by comparison to a reference dataset (41).

## Results

### Overview of FANS workflow

We present a series of optimised FANS protocols enabling the isolation of multiple neural cell-type–enriched nuclear populations from frozen human and mouse cortex (Figure 1). These methods are compatible with both pre-and post-natal tissue and require only small amounts of starting material. Across all applications, the FANS workflow consistently yielded nuclei of sufficient purity and quantity for downstream transcriptomic, DNA modification, chromatin accessibility, and histone profiling assays. The results presented below illustrate the performance of the protocol and validate its suitability for high-quality, multiomic analysis of human and mouse brain tissue.

### FANS enables isolation of distinct nuclei populations for multiomic analysis

FANS was employed to isolate distinct nuclear populations from human and mouse cortical tissue, enabling their use in multiple downstream assays. The gating strategy was designed hierarchically so that each sorted population represents a defined subpopulation (“daughter”) within the total nuclei gate. This strategy ensured that proportions were normalised relative to the entire nuclei pool, with consistent recovery across samples. On average, neurons comprised 34.5% ± 13.5, oligodendrocytes 46.7% ± 15.9, microglia 3.3% ± 1.8 and astrocytes 3.2% ± 3.15 of the total sorted events. These distributions confirm that neurons and oligodendrocytes are the predominant nuclear populations in the human cortex, with microglia and astrocytes present in lower relative abundance.

From approximately 300 mg of postnatal human cortical tissue, the FANS protocol routinely enables the recovery of 50,000 nuclei for DNA methylation analysis, 300,000 nuclei for nuclear RNA extraction, 100,000 nuclei for chromatin accessibility assays (ATAC-seq), and 100,000 nuclei for histone modification profiling (e.g., H3K27ac CUT&Tag) for each tested cell-type. In mouse cortex, the protocol robustly yields 60,000 NeuN⁺ (neuron-enriched) nuclei, 5,000 PU.1⁺ (microglia-enriched) nuclei, and 20,000 (NeuN⁻/PU.1⁻; oligodendrocyte-enriched) nuclei from ∼100 mg of frozen tissue. Nuclei recovery from human and mouse cortex showed variability across samples, largely reflecting inter-individual differences and heterogeneity between cortical subregions. Factors such as dissection accuracy, lipid content, and developmental stage can influence overall yield and population representation.

### Validation of antibody specificity and sorting purity in postnatal human cortex

To validate antibody specificity and assess sorting purity, microscopy imaging was performed on immunolabeled nuclei before FANS (**Figure S4 & Figure S5**). These images confirmed that the selected antibodies specifically label distinct nuclear populations, consistent with the expected cellular heterogeneity of the postnatal cortex. The presence of a diverse mixture of labelled nuclei supports the suitability of the chosen markers for distinguishing major brain cell types. Sorting purity was further assessed through reanalysis of sorted populations. **Figure S6** presents a representative example of a NeuN⁺ population subjected to re-sorting, demonstrating high purity and minimal contamination. This analysis confirms the reliability, specificity, and reproducibility of the FANS protocol in isolating well-defined nuclear subpopulations from complex brain tissue

### Targeted and genome-wide gene expression analyses confirm cell-type identity

To confirm the enrichment of cell-type-specific transcripts within FANS-sorted nuclei populations, targeted qPCR gene expression analysis was performed on nuclear RNA extracted from isolated nuclei populations. Transcripts corresponding to canonical marker genes for the major brain cell types were quantified, including *ENO2* and *RBFOX3* (neurons), *OLIG2* (oligodendrocytes), *GFAP* (astrocytes), and *CD68* (microglia). Gene expression levels were normalised to five reference genes previously validated for stable expression in human brain tissue (42). As expected, the expression of *RBFOX3* - a marker of mature neurons - was highly enriched in the NeuN⁺ nuclei fraction (**Figure S7a**), with minimal expression observed in both the SOX10⁺ and NeuN⁻/SOX10⁻ fractions. In contrast, *OLIG2* expression was strongly enriched in the SOX10⁺ nuclei, and nearly undetectable in the other two populations, consistent with its known specificity for oligodendrocytes. The NeuN⁻/SOX10⁻ (double-negative or antibody-negative) population displayed elevated expression of both *GFAP* and *CD68*, markers for astrocytes and microglia, respectively, suggesting that this fraction is enriched for non-neuronal, non-oligodendrocyte glial cells (astrocytes and microglia). To complement the targeted analysis, bulk RNA-seq was performed on the NeuN⁺ and SOX10⁺ nuclei populations. RNA-seq libraries were generated from 300,00 nuclei per sample, yielding an average of 85 million reads per library. The transcriptomic profiles obtained confirmed the cell-type-specific enrichment patterns observed by qPCR (**Figure S7b**). Together, these data confirm the successful isolation and molecular characterisation of major brain cell types using FANS and provide transcriptional validation of population identity across both targeted and genome-wide expression modalities.

### Single-nucleus RNA-seq validates confirms distinct transcriptional profiles of FANS-sorted nuclei population

To validate the utility of FANS-sorted nuclei for single-nucleus transcriptomics, we performed snRNA-seq using both Parse Biosciences (80,151 nuclei) and 10x Genomics Chromium (9,448 nuclei). In both datasets, nuclei clustered primarily by FANS-enriched population rather than individual donor (Figure 2h and **Figure S8**), demonstrating minimal batch effects and strong biological signal. Parse data displayed distinct transcriptional clusters corresponding to neurons, oligodendrocytes, microglia, and mixed glia (Figure 2g**–l**). Marker gene expression patterns within these clusters matched the expected identities of each FANS-isolated population (**Figure S9**). Similarly, 10x datasets confirmed high expression of *SYT1* in NeuN⁺ nuclei, *OLIG2* and *PLP1* in SOX10⁺ nuclei, and *PTPRC* and other microglial markers in IRF8⁺ nuclei (**Figure S8**). These analyses highlight the compatibility of FANS with single-nucleus platforms and confirm the high specificity of enrichment.

### DNA methylation profiling confirms cell-type-specific patterns

Genome-wide DNA methylation (DNAm) profiling was performed on FANS-sorted nuclear populations from human postnatal cortex using the Illumina EPIC v2 array. After stringent QC and filtering, a total of 889,069 high-quality DNA methylation sites remained for downstream analysis. The final dataset used to validate the FANS protocol included DNA methylation profiles from 47 individuals (14 female, 33 male; age range: 6–91 years), each with up to four nuclear fractions profiled: NeuN⁺ (n = 44), Sox10⁺ (n = 44), IRF8⁺ (n = 42), and triple-negative (TN) nuclei (n = 45). To validate cell-type purity and the effectiveness of the sorting strategy, we applied a neural cell-type deconvolution model (25) to the EPIC array data confirming very high purity for all sorted fractions (>90%), consistent with the transcriptomic validation. (**Figure S10**). Unsupervised hierarchical clustering and heatmap visualisation of the 1,000 most variable DNA methylation sites across samples further demonstrated robust cell-type-specific methylation signatures (Figure 2f). Samples clustered by nuclear fraction, with clear separation between neurons, oligodendrocytes, microglia, and other glia. Notably, CpG sites within known marker genes for each cell type exhibited expected, cell-type-specific methylation patterns (**Figure S11**). For additional examples of DNA methylation-based clustering in postnatal human please refer to Franklin et al. (2025) (43). Clustering of mouse FANS-sorted cortical samples is shown in Supplementary **Figure S12**. Finally, ONT genomic sequencing data from DNA isolated from three nuclei populations (NeuN^+^, SOX10^+^ and IRF8^+^), isolated via FANS from three postnatal human PFC samples, revealed distinct cell-type differences in both 5mC and 5hmC, as illustrated in Figure 3 and **Figure S13**, demonstrating that the protocol is suitable for long-read, base-resolution epigenomic assays.

**Figure 3.**
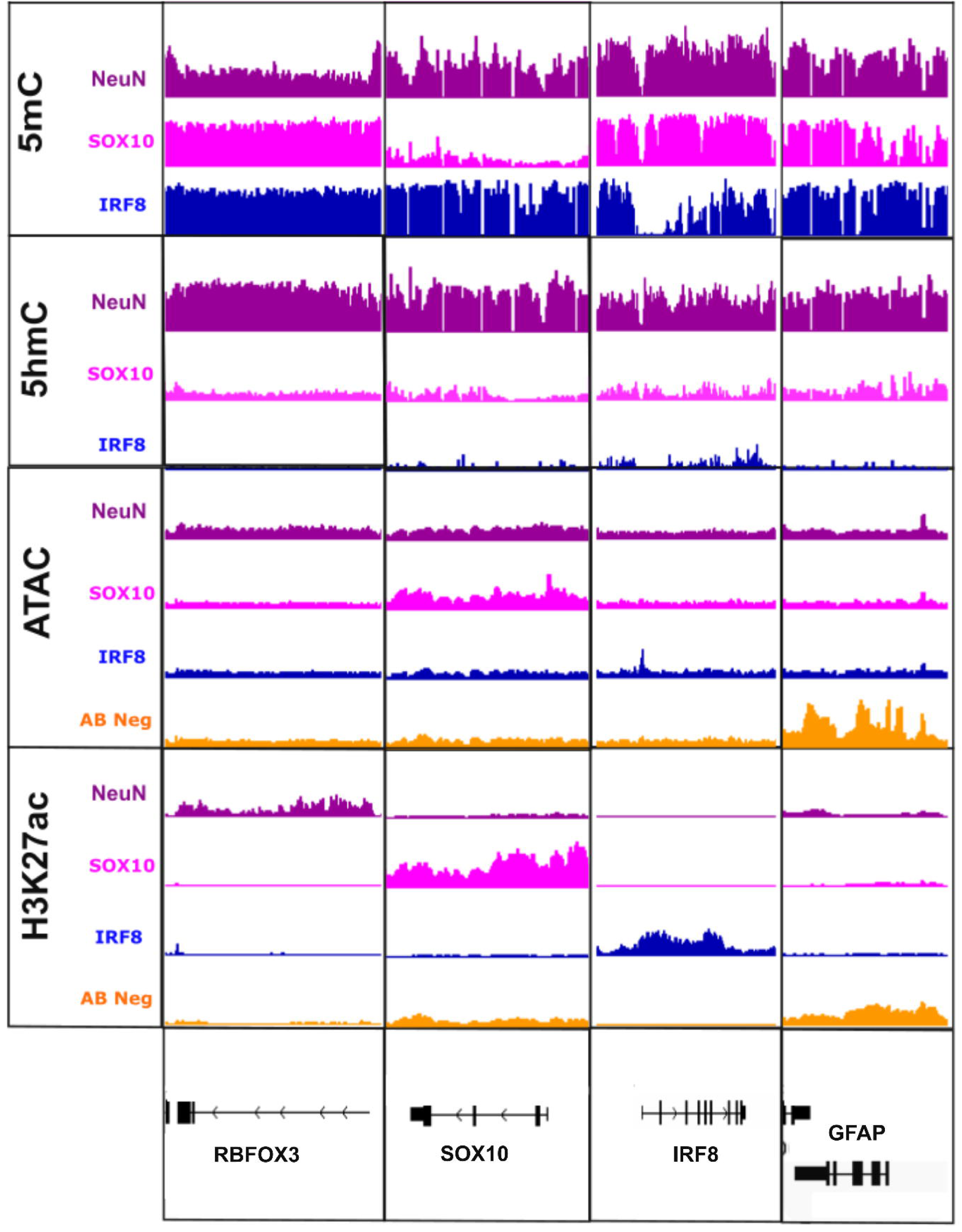
Integrated multi-omic profiling reveals consistent cell-type–specific signatures in FANS-isolated nuclei. Genome-wide DNA modification signals from Oxford Nanopore Technologies (ONT) sequencing, chromatin accessibility profiles from ATAC-seq, and H3K27ac enrichment from CUT&Tag were visualised using IGV across representative marker genes for each major brain cell type. Across all three modalities, nuclei isolated by FANS show distinct and internally consistent epigenomic patterns characteristic of neurons (NeuNL), oligodendrocytes (SOX10L), microglia (IRF8L), and astrocyte-enriched triple-antibody negative (ABL) populations. These concordant profiles across diverse genomic assays demonstrate the specificity, robustness, and effectiveness of the FANS-based nuclei isolation workflow.

### ATAC-seq analysis reveals distinct chromatin landscapes in sorted nuclei populations

After stringent quality control and filtering steps, we obtained high-quality ATAC-seq data from 110 NeuN⁺, 119 SOX10⁺, 24 IRF8⁺ and 27 triple-negative (NeuN⁻/SOX10⁻/IRF8⁻; referred to as TN) nuclei populations isolated from human postnatal cortex. Fragment size distributions showed the expected periodicity for nucleosome-free and mononucleosomal fragments (**Figures S14 and S15**), confirming high-quality library preparation. Peak calling identified hundreds of thousands of accessible regions per population, with distinctive and reproducible accessibility profiles. Unsupervised sample clustering (**Figure S16**) revealed strong separation between neuronal, oligodendrocyte, microglial, and TN nuclei, confirming cell-type-specific chromatin accessibility patterns.

### H3K27ac CUT&Tag confirms cell-type-specific enhancer landscapes

CUT&Tag profiling of H3K27ac produced 410 high-quality libraries from four major FANS-sorted populations obtained from human postnatal cortex. After filtering, each population exhibited a distinct enhancer landscape, with strong within-cell-type correlations (0.89-0.96) and markedly lower correlations between different cell types (0.05-0.43) (**Figure S17**).

Principal component analysis confirmed that the first three components captured variation between oligodendrocyte (SOX10^+^), microglial (IRF8^+^), neuronal (NeuN^+^), and TN nuclei (data not shown). These results demonstrate that FANS produces nuclei suitable for sensitive chromatin profiling of histone modifications.

### Integrated multiomic validation demonstrates cell-type–specific genomic signatures in FANS-Isolated populations

Integration of DNA methylation, chromatin accessibility (ATAC-seq), H3K27ac CUT&Tag, and ONT modification data at canonical marker genes (Figure 3) revealed coherent and highly cell-type-specific epigenomic signatures. Neurons, oligodendrocytes, microglia, and TN glia each displayed characteristic methylation, chromatin accessibility, and acetylation patterns, consistent across individuals and assays. These findings demonstrate that FANS yields nuclei that capture the authentic molecular identity of each major brain cell type, enabling parallel multiomic profiling from the same tissue sample.

## Discussion

The protocols described here provide a framework for generating high-quality, cell-type–specific multiomic data from frozen human and mouse brain tissue. These methods are designed to support downstream analyses across a range of omics approaches including transcriptomics, DNA and histone modification profiling, and chromatin accessibility assays. By maximising the informational yield from limited and often precious cortical samples, our protocols provide a robust framework for multidimensional molecular profiling. This integrated approach enhances the resolution at which disease-associated variation can be investigated in both human and model systems, thereby advancing our understanding of neurodevelopment and neurodegeneration.

Through methodological refinement of fluorescence-activated nuclei sorting (FANS), we reliably isolated neurons (NeuN⁺), oligodendrocytes (SOX10⁺), microglia (IRF8⁺), and astrocyte-enriched (antibody-negative) nuclei in postnatal human cortex, as well as equivalent populations in fetal and mouse cortex. These protocols build upon earlier methods for neuronal nuclei isolation for ChIP-seq (44) and snRNA-seq (45)but introduce significant improvements to capture a broader range of cell types and facilitate multiomic profiling. We extend these approaches to reliably isolate oligodendrocytes and microglia, using SOX10 and IRF8, respectively, as nuclear markers. Although previous studies have also used alternative markers such as OLIG2 and PU.1 and other cell types (46) our protocols are distinct in their optimisation for multi-omic applications, enabling transcriptomic, epigenomic, and chromatin accessibility profiling from the same nuclear preparations. A major advance demonstrated here is the ability to obtain four complementary molecular datasets (RNA-seq, DNA modification profiling (array and ONT), ATAC-seq, and histone modification mapping via CUT&Tag) from a single ∼300 mg cortical tissue sample. This efficiency is particularly valuable in the context of human post-mortem brain research, where tissue availability is limited and finite. The consistency of cell-type-specific signatures across all modalities provides strong validation of the specificity and purity of FANS-isolated fractions. In addition, we have optimised our FANS protocol to include markers for other distinct neural cell types, such as motor neurons (BCL6), GABAergic neurons (SOX6), and astrocytes (LHX2). Highlighting that this strategy can be adapted to a wide range of cell types, including unique or low-abundance populations, as long as an appropriate antibody marker is available.

Despite the strengths of our protocol, several limitations warrant consideration. First, the use of post-mortem tissue inherently introduces variables related to tissue quality, post-mortem interval, and donor variability. Second, although we confirm the positive purification of nuclei from different cell types, the developed protocols cannot purify nuclei from more specific cellular subtypes. For example, NeuN is a widely used and reliable pan-neuronal marker; however, it has important limitations that constrain its ability to distinguish neuronal subtypes. NeuN is not uniformly expressed across all neuronal populations - in particular, it is absent or weakly expressed in certain classes of neurons such as cerebellar Purkinje cells, mitral cells of the olfactory bulb, and some retinal neurons (47). As a consequence, NeuN-based sorting enriches for the majority of cortical neurons but does not capture the full diversity of neuronal lineages, and it cannot differentiate between excitatory and inhibitory neurons or between distinct projection neuron subtypes. Newer cytometry platforms with expanded fluorophore capabilities will enable the capture of more distinct cell populations, such as separating excitatory versus inhibitory neurons or differentiating reactive from homeostatic microglia.

Expanding the panel of nuclear markers and developing novel, highly specific nuclear antibodies will be essential for resolving cellular subpopulations and disease-relevant cellular states. This will ultimately enhance the power of single-nucleus omics to interrogate brain development, plasticity, and pathology at unprecedented resolution.

## Conclusion

The protocols presented here provide a reliable, scalable, and highly adaptable strategy for cell-type–specific nuclei isolation and multiomic profiling from frozen brain tissue. By enabling comprehensive molecular characterisation of multiple neural populations from the same sample, these methods maximise the value of limited post-mortem resources and open new avenues for mechanistic studies of human brain function and pathology. They represent a significant methodological advance for the field and will support future efforts to map regulatory architectures, model disease mechanisms, and identify cell-type–specific therapeutic targets across the human lifespan.

## Supporting information

Protocol 1

Protocol 2

Protocol 3

## Acknowledgements

These data were generated as part of Medical Research Council grant K013807 to JM and Alzheimer’s Research UK (ARUK) grant ARUK-PPG2018A-010 to E.L.D. E.H, J.M, E.L.D and S.M were supported by Medical Research Council (MRC) grants K013807 and W004984 (awarded to J.M.). E.H is supported by an Engineering and Physical Sciences Research Council Fellowship EP/V052527/1. Data analysis was undertaken using high-performance computing supported by a Medical Research Council (MRC) Clinical Infrastructure award (M008924) to J.M. This study was supported by the National Institute for Health and Care Research Exeter Biomedical Research Centre. The views expressed are those of the author(s) and not necessarily those of the NIHR or the Department of Health and Social Care. We acknowledge the supply of samples from Human Developmental Biology Resource at the IGM, Newcastle upon Tyne; The Guy’s and St Thomas’ Research Biobanks; The Institute of Child Health, University College London, London; Kings College London in association with the MRC London Brain Bank for Neurodegenerative Diseases and funders of the Brain Bank; The South London and Maudsley NHS Foundation Trust; The London Neurodegenerative Diseases Brain Bank, Brains for Dementia Research; JJ Peters VA Medical Center; the Oxford Brain Bank, supported by the Medical Research Council (MRC), the NIHR Oxford Biomedical Research Centre and the Brains for Dementia Research programme, jointly funded by Alzheimer’s Research UK and Alzheimer’s Society.

