## Supplementary material for "Optimised fluorescence-activated nuclei sorting for epigenomic analysis of cortical cell types": Protocol 1

Nov 27, 2024 Version 2

### Fluorescence-activated nuclei sorting (FANS) on human post-mortem cortex tissue enabling the isolation of distinct neural cell populations for multiple omic profiling V.2

DOI
dx.doi.org/10.17504/protocols.io.36wgq4965vk5/v2

Stefania S Policicchio[1], Barry Chioza[1], Jonathan P Davies[1], Joe Burrage[1], Jonathan Mill[1], Emma L Dempster[2]

[1]University of Exeter, Exeter, UK; [2]University of Exeter

Complex Disease Epige...

Stefania S Policicchio
University of Exeter Medical School

OPEN ACCESS

**DOI:** https://dx.doi.org/10.17504/protocols.io.36wgq4965vk5/v2

**External link:** https://doi.org/10.1038/s41467-022-33394-7

**Protocol Citation:** Stefania S Policicchio, Barry Chioza, Jonathan P Davies, Joe Burrage, Jonathan Mill, Emma L Dempster 2024. Fluorescence-activated nuclei sorting (FANS) on human post-mortem cortex tissue enabling the isolation of distinct neural cell populations for multiple omic profiling. **protocols.io** **https://dx.doi.org/10.17504/protocols.io.36wgq4965vk5/v2**Version created by **Barry Chioza**

**Manuscript citation:**
Shireby G, Dempster EL, Policicchio S, Smith RG, Pishva E, Chioza B, Davies JP, Burrage J, Lunnon K, Vellame DS, Love S, Thomas A, Brookes K, Morgan K, Francis P, Hannon E, Mill J, DNA methylation signatures of Alzheimer's disease neuropathology in the cortex are primarily driven by variation in non-neuronal cell-types. Nature Communications doi: **10.1038/s41467-022-33394-7**

**License:** This is an open access protocol distributed under the terms of the **Creative Commons Attribution License**, which permits unrestricted use, distribution, and reproduction in any medium, provided the original author and source are credited

**Protocol status:** Working
**We use this protocol and it's working**

**Created:** October 04, 2023

**Last Modified:** November 27, 2024

**Protocol Integer ID:** 88785

**Keywords:** FANS, post-mortem brain, nuclei, flow cytometry, anti-SOX10, anti-IRF8, anti-NeuN, nuclei sorting, nuclei from multiple different human brain cell, cellular composition in regulatory genomic study, multiple different human brain cell, heterogeneous mix of different neural cell type, distinct neural cell population, isolation of distinct neural cell population, epigenetic process, regulatory genomic study, other glial origin nuclei, different neural cell type, open chromatin analysis, genomic data, genome, gene regulation, specific cell, specific patterns of gene regulation, oligodendrocyte, transcriptional variation in health, cellular composition, amount of genomic data, determining cell type, tissue sample, transcriptional variation, current analyses of the human brain, cell type, gene expression, human brain, dna, populations of specific cell, microglia, cell, multiple omic profiling

**Funders Acknowledgements:**
**MRC Research Grant**
Grant ID: MR/R005176/1
**Alzheimer's Research UK (ARUK)**
Grant ID: ARUK-PPG2018A-010

#### Abstract

Increased understanding of the functional complexity of the genome has led to growing recognition about the role of epigenetic/transcriptional variation in health and disease. Current analyses of the human brain, however, are limited by the use of "bulk" tissue, comprising a heterogeneous mix of different neural cell types.
As epigenetic processes play a critical role in determining cell type-specific patterns of gene regulation it is important to consider cellular composition in regulatory genomic studies of human post-mortem tissue, and there is a need for methods to purify populations of specific cell-types. Furthermore, the valuable nature of human post-mortem tissue means it is important to use methods that maximize the amount of genomic data generated on each sample.
This protocol describes a method that uses fluorescence-activated nuclei sorting (FANS) to isolate and profile nuclei from multiple different human brain cell-types from frozen post-mortem tissue.
This protocol can be used to robustly purify populations of neuronal (NeuN$^{+ve}$), oligodendrocytes (SOX10$^{+ve}$), microglia (IRF8$^{+ve}$) and other glial origin nuclei (NeuN$^{-ve}$/SOX10$^{-ve}$/IRF8$^{-ve}$) from adult post-mortem frozen brain, with each tissue sample yielding purified populations of nuclei amenable to simultaneous analysis of i) DNA modifications (via bisulfite sequencing / array), ii) histone modifications (CUT&Tag), iii) open chromatin analysis (ATAC-seq), and iv) gene expression (snRNA-seq).

#### Guidelines

Prior to attempting this protocol please obtain the approval of your Institutional Review Board (IRB) or equivalent ethics committee(s).

#### Materials

| | A | B | C |
|---|---|---|---|
| | | **Supplier** | **Catalogue No.** |
| | **BD FACSAria™ III Cell Sorter** | BD Biosciences | 648282-23 |
| | **Sorvall WX 80+ Ultracentrifuge** | Thermo Scientific™ | 75000080 |
| | **7mL Dounce Tissue Grinder** | DWK Life Sciences | 357542 |
| | **PA Thin-walled ultracentrifuge tubes** | Thermo Scientific | 10533934 |

***Table 1:*** Specifications of the equipment required for FANS protocol

| | A | B | C |
|---|---|---|---|
| | | **Supplier** | **Catalogue No.** |
| | **D-Sucrose (Molecular Biology)** | Fisher Scientific | 10638403 |
| | **Calcium chloride (CaCl2) anhydrous, granular** | Sigma-Aldrich | C1016-100G |
| | **Magnesium acetate ( Mg(Ace)2), 1M aq. soln** | Alfa Aesar | J60041 |
| | **UltraPure 0.5M EDTA, pH 8.0** | Invitrogen | 15575020 |
| | **Thermo Scientific 1M Tris-HCl Buffer, pH 8.0** | Fisher Scientific | 15568025 |
| | **1,4-Dithiothreitol (DTT) - crystalline powder** | Sigma-Aldrich | 3483-12-3 |
| | **Triton X-100** | Sigma-Aldrich | T9284 |
| | **UltraPure DNase/RNase-Free Distilled Water (ddH2O)** | Fisher Scientific | 12060346 |
| | **Bovine Serum Albumin (BSA)** | Sigma-Aldrich | A9647-500G |
| | **PBS Phosphate-Buffered Saline (10X) pH 7.4** | Fisher Scientific | 10722497 |
| | **Thermo Scientific™ RiboLock RNase Inhibitor (40 U/µL)** | Fisher Scientific | 10389109 |
| | **TRIzol LS Reagent** | Invitrogen | 11588616 |
| | **BAMBANKER serum free cell freezing medium** | BioCat GmbH | BB03-NP |
| | **BD FACSDiva CS&T Research Beads** | BD Biosciences | 655051 |
| | **BD FACS Accudrop Beads** | BD Biosciences | 345249 |

| A | B | C |
|---|---|---|
| **BD FACSFlow Sheath Fluid 20L** | BD Biosciences | 342003 |
| **BD FACS Clean Solution** | BD Biosciences | 15875858 |
| **BD FACS Rinse Solution** | BD Biosciences | 340346 |

***Table 2*:** Specification of reagents required for FANS protocol

| A | B | C |
|---|---|---|
| **Lysis Buffer (LB)** | | |
| | **Stock** | **Amount** |
| 0.32M Sucrose | | 5.47 g |
| 5mM CaCl2 | 1M | 250 µL |
| 3mM Mg(Ace)2 | 1M | 150 µL |
| 0.1mM EDTA | 0.5M | 10 µL |
| 10mM Tris-HCl, pH 8 | 1M | 500 µL |
| 1mM DTT | 3M | 17 µL |
| 0.1% Triton X-100 | | 50 µL |
| *Optional: RiboLock RNase Inhibitor 0.2U/µL* | *40U/ µL* | *5 µL / 1mL* |
| Adjust with ddH2O to | | 50 mL |
| **1.8M Sucrose Solution (SS)** | | |
| 1.8M Sucrose | | 30.78 g |
| 3mM Mg(Ace)2 | 1M | 150 µL |
| 1mM DTT | 3M | 17 µL |
| 10mM Tris-HCl, pH8 | 1M | 500 µL |
| Adjust with ddH2O to | | 50 mL |
| **5% BSA Solution (BB)** | | |

| | A | B | C |
|---|---|---|---|
| | BSA | | 200 mg |
| | 1x PBS | | 4 mL |
| | **Staining Buffer (SB)** | | |
| | 0.5% BSA | 5% BSA Solution (BB) | 400 µL |
| | 10X PBS | | 400 µL |
| | *Optional: RiboLock RNase Inhibitor 0.2U/µL* | *40U/ µL* | *5 µL / 1mL* |
| | Adjust with ddH2O to | | 4 mL |

***Table 3*** : Recipes for buffers and solutions required

| | A | B |
|---|---|---|
| | **Supplier** | Thermo Scientific™ |
| | **Model** | Sorvall™ WX 80+ |
| | **Rotor** | TH-641 |
| | **Speed** | 25,200 RPM / 108670.8 x g |
| | **Acceleration** | 9 |
| | **Deceleration** | 5 |
| | **Temperature** | 4°C |

***Table 4:*** *Ultracentrifuge specification and conditions*

| | A | B | C | D | E |
|---|---|---|---|---|---|
| | **Antibody** | **Preconjugated** | **Supplier** | **Cat No** | **Dilution** |
| | Hoechst 33342 | -- | Abcam | ab228551 | 1:500 |
| | Anti-SOX10 | to NL557 | R&D systems | NL2864R | 1:20 |
| | Anti-NeuN | to Alexa Fluor488 | Millipore | MAB377X | 1:1000 |
| | Anti-IRF8 | to APC | Invitrogen | 17-9852-82 | 1:150 |

***Table 5***: List of antibodies required for FANS protocol

#### Protocol materials

TRIzol LS **Fisher Scientific Catalog** #11588616

BAMBANKER **BioCat GmbH Catalog** #BB03-NP

#### Troubleshooting

#### Nuclear prep for FACS separation (using SOX10, IRF8, NeuN and Hoechst)

1 The protocol below yields at least 1,000,000 NeuN +ve, 1,000,000 SOX10 +ve, 400,000 IRF8 +ve (when the population is present) and 200,000 triple negative (NeuN-ve/SOX10-ve/IRF8-ve) nuclei per 500 mg of frozen human post-mortem cortex tissue. Recovery might vary from sample to sample due to high inter-sample variability (brain collection, cortex sub-areas, fat content of tissue sectioned, and white to grey matter ratio)

Refer to Materials-***Table 1*** for details about the equipment required and to Materials-***Table 2*** for specifications of reagents required.

1.1 ***Solution and buffer preps***

- *Lysis Buffer (LB)*
- *Sucrose Solution (SS)*
- *Staining Buffer (SB)*

Solutions should be kept at 4 °C or On ice. Refer to Materials-***Table 3*** for recipes of solutions and buffers.

Note

**NOTE 1** – LB and SS can be prepared a week in advance, with DTT added on the day of use. Solutions should be stored at 4 °C once made.

Note

**NOTE 2** – SB should be prepared fresh each day.

Note

**NOTE 3**- Samples are homogenised as bulk tissue using a 7 mL Dounce homogeniser and then equally divided into three ultracentrifuge tubes.

1.2 ***Nuclei isolation*** 1h 20m

1. Pre-cool the ultracentrifuge to 4 °C 00:30:00 before starting this stage of the protocol.
2. All buffers and the Dounce homogenisers should be pre-cooled on ice.

3. Add DTT ( [M] 1 millimolar (mM) final concentration ) to the SS and LB (i.e. Amount 17 µL 3M DTT per 50 mL of SS/LB)
4. Transfer 3 mL LB to the homogeniser per 500 mg human brain tissue.
5. Add the dissected tissue sample into the homogeniser.
6. Wait 00:05:00 before douncing the tissue to allow the sample to defrost.

To reduce heat caused by friction, the Dounce homogenisation step should be performed on ice with gentle strokes, and care should be taken to avoid foaming.

Note

**NOTE 1** – Using the "TIGHT" pestle helps reduce the number of strokes required to reach full tissue disruption.

Note

**NOTE 2** - The number of strokes required to fully homogenise the tissue may vary between samples due to heterogeneity in cellular composition, lipid content, and the amount of connective tissue.

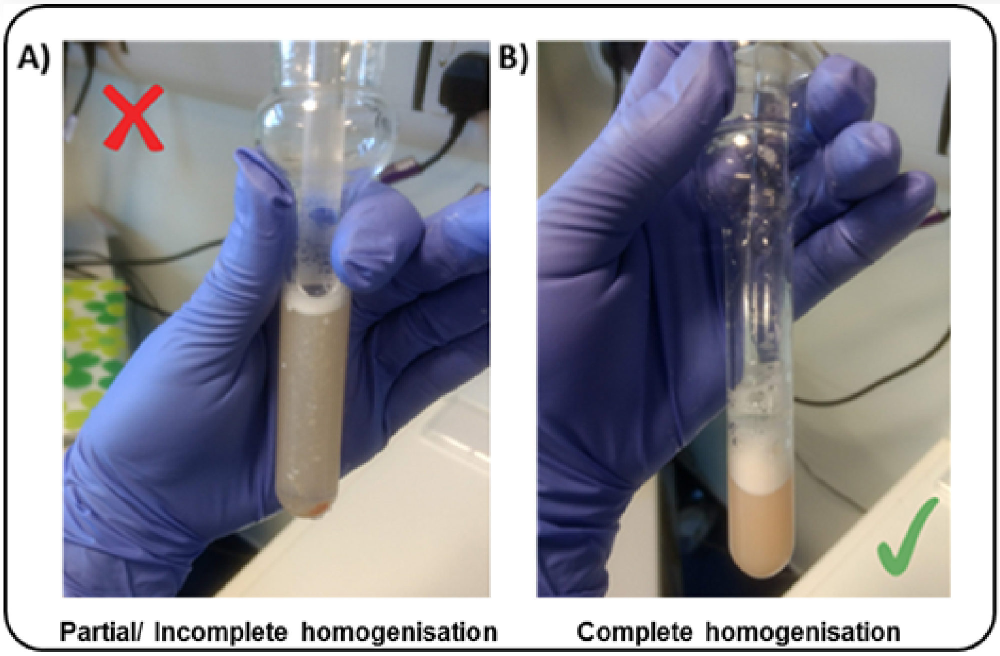

***Figure 1*** Example of brain tissue sample **A)** only partially homogenised **B)** complete homogenisation.

7. Transfer 8 mL SS to PA thin-walled ultracentrifuge tubes.
8. Carefully overlay with tissue homogenate ( 1 mL per tube) - using a P1000 pipette, releasing slowly down the side of the tube.

9. Overlay with another 1 mL LB – do not worry about disrupting the homogenate phase
10 .Balance opposite tubes by weight with LB using a fine microbalance.
11. Perform ultracentrifugation for 00:45:00 (see ***Table 4*** for centrifuge specification and conditions).

1.3 ***After Ultracentrifugation step*** 20m

1. Pour off any remaining supernatant, taking care not to dislodge the pellet (90-degree inclination of the tube). If the pellet is hard to see, it is okay to leave solution in the ultracentrifuge tube.
2. Re-suspend pellet in 1 mL SB and gently pipette up and down.
3. Let samples sit on On ice for at least 00:15:00

**Blocking step**

4. Transfer volume into 2 mL Lo-Bind Eppendorf tubes.
5. Rinse out ultracentrifuge tubes in order to maximise nuclei collection by adding SB to each ultracentrifuge tube ( 1 mL ) pipetting up and down several times and transferring to the new 2 mL tubes.

**Washing step:**

6. Centifuge tubes at 1000 x g for 00:05:00 , 4 °C
7. Discard supernatant (pipetting off gently).
8. Re-suspend each nuclei pellet in fresh SB ( 500 µL )

If the sample was split then pool together pellets from the same sample (Final Volume = 1.5 mL )

9. Add DNA dye (Hoechst, 1:1000) and mix thoroughly via inversion.
10. Pipette out 200 µL of nuclei solution for the Unstained Control (Hoechst dye only) and transfer to a new 2 mL tube
11. Bring the volume up to 1 mL for the Unstained tube with fresh SB.
12. Replace the volume taken from the Stained" tube with 200 µL of fresh SB (Final Volume 1.5 mL ).

1.4 ***Immunostaining***

1. Add the following three antibodies (Ab) to Stained tube (1.5ml):

- SOX10 pre-conjugated antibody (1:20 dilution) – [ 75 µL Ab]
- NeuN Alexa488 (1:1000) – [ 1.5 µL Ab ]
- IRF8 pre-conjugated antibody (1:150) - [ 10 µL Ab]

Refer to ***Table 5*** for specifications of the antibodies used

2. Incubate tubes for at least 01:30:00 on the rotor mixer (speed=11) at 4 °C , keeping the tubes in the dark
3. Washing step: 1000 x g for 00:05:00 , 4 °C (both "Stained" and "Unstained" tubes)
4. Discard supernatant (by aspiration)
5. Re-suspend in fresh SB ( 0.5 mL for the Unstained tube, -2 mL for the Stained tube - depending on pellet size)

#### Fluorescence-Activated Nuclei Sorting (FANS)

2 For machine start-up, CST and Accudrop calibrations refer to **BD FACSAria III User's Guide** for guidance and troubleshooting. The following instructions describe FANS using BD FACSAria III. Other FACS platforms can be used but might require modifications to the protocol.

2.1 ***General Gating Parameters***

For each sample, load stained and unstained tubes individually for data acquisition. A preliminary qualitative analysis of the data acquired is essential to select the appropriate gating strategy to maximize the nuclei capture while excluding unnecessary debris and to ensure optimal signal/noise ratio.

*Gating Parameters (X-axis:Y-axis):*

FSC-A:SSC-A (Size, cell granularity or internal complexity)
SSC-W:SSC-A (to gate out doublets)
FSC-A:DAPI-A (to gate the single nuclei population)
DAPI-A:FITC-A (to gate NeuN stained nuclei)
DAPI-A:PE-A (to gate SOX10 stained nuclei)
DAPI-A:APC-A (to gate IRF8 stained nuclei)
FITC-A: PE-A (to visualize the distribution of triple staining)

Note

The SOX10$^{+ve}$ population is gated as a “daughter” population from the NeuN$^{-ve}$ fraction.
The IRF8$^{+ve}$ population is gated as a “daughter” population of the SOX10$^{-ve}$ fraction.
Refer to **Figure 2** for a visualization of the gating strategy.

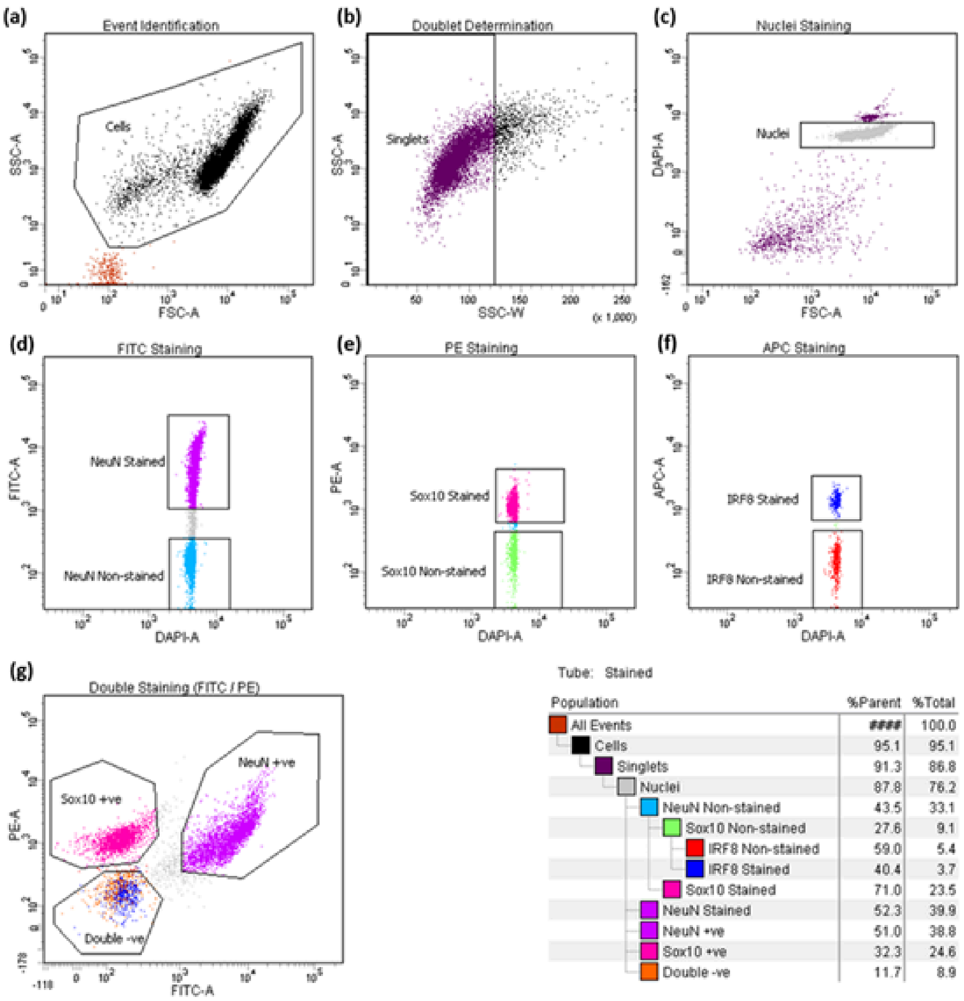

**Figure 2.** FANS gating strategy.
**(a)** Particles smaller than nuclei (black dots) were eliminated with an area plot of forward-scatter (FSC-A) versus side-scatter (SSC-A), with gating for nuclei-sized particles inside the gate (box).
**(b)** Plots of height versus width in the side scatter channel are used for doublet discrimination with gating to exclude aggregates of two or more nuclei.
**(c)** Doublet discrimination gating was used to isolate nuclei determined by subgating on Hoechst 33342.
Subsequent scatterplots discerning :
**(d)** NeuN-Alexa Fluor488–conjugated antibody staining (purple)
**(e)** SOX10-NL577-stained nuclei (dark pink)
**(f)** IRF8-APC stained nuclei (dark blue)
**(g)** the distribution of the three main nuclei subpopulations identified through triple staining strategy (NeuN$^{+ve}$, neurons; SOX10$^{+ve,}$ oligodendrocytes; double$^{-ve}$, glia).
The resultant hierarchical colour key ensures that only nuclei that are positive or negative for staining with the NeuN and/or SOX10/IRF8 antibody are passed through each gating condition.

2.2 ***Data recording settings***

In line with the experiment design, FSC, SSC, DAPI, FITC, APC and PE are the parameters for which voltage values may need to be slightly adjusted due to experiment/ inter-sample variability.

It is advisable to set the threshold value between 200 and 500 during data recording. Moreover, in the acquisition dashboard tab, we recommend setting **Events to Record** ≤3000, **Event to display** ≤1000 and **Flow Rate** = 1.0 (1,000 events per second) in order to increase the accuracy of signal detection.

The flow rate can be increased during sample collection to reduce the sort speed (ideally max events per second =1,500 for a 100-micron nozzle). However higher flow rates impact the data resolution and accuracy of events detection, and subsequent sorting of cellular fractions (see **BD FACSAria III User's Guide** for details).

During analysis, recorded data is displayed in plots, while gates are used to define populations of interest for selection. **Figure 3** shows a representative example of the two most common outcomes we often observe.

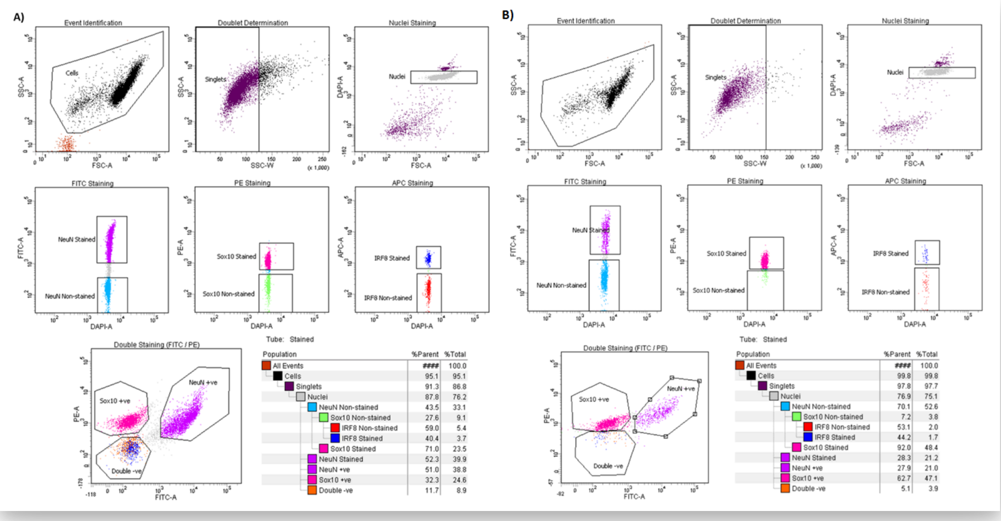

**Figure 3**. Representative example of inter-individual variability.

The data shown here are derived from two different control prefrontal cortex specimens of comparable age, gender, and brain collection which were processed in parallel following the same procedure.

**A)** Optimal sample separation and abundant IRF8$^{+ve}$ fraction vs **B)** poor sample separation and completely missing a double negative population or an IRF8$^{+ve}$ population.

2.3 ***Sample Collection*** 5m

1. DNA LoBind Tubes (Eppendorf, Cat No:30108051) are required to collect nuclei (to maximize sample recovery of nucleic acids by significantly reducing sample–to–surface binding).
2. Collected fractions can be used directly for downstream applications (e.g. DNA/RNA extraction, ATAC-seq, etc) or stored in -80 °C freezer.
3. If you are collecting for DNA or RNA, as soon as the number of events desired is reached, transfer the tubes On ice , do not hold them at Room temperature

Note

During collection, it is crucial to regularly pause the sorting to mix the two phases in order to preserve the integrity of resulting RNA preparations.

Note

For RNA extraction, LoBind Tubes should each contain 1 mL of pre-chilled TRIzol LS **Fisher Scientific Catalog** #11588616 prior to sorting.

4. Keep samples On ice for the entire duration of the sorting.
5. Lightly vortex sample tubes to make the mixture homogeneous (not clumped) before loading the tube into the FACS chamber.
6. Load the **UNSTAINED** control tube into the chamber first and proceed with nuclei collection.
7. Proceed by collecting **STAINED** tube by simultaneously sorting for $NeuN^{+ve}$, $SOX10^{+ve}$, $IRF8^{+ve}$ or Triple Negative.

Note

At least 1 µg genomic DNA at least is expected from 500 mg tissue. For optimal recovery of high-quality genomic DNA from FANS sorted nuclei we recommend this **extraction protocol**

Note

The $IRF8^{+ve}$ population may not be detectable in every brain sample processed (high inter-individual variability); when it is, it represents between 5-10% of the total sample, therefore yielding often insufficient material for multiple assays.

8. For long term storage of collected nuclei 1000 x g , 4 °C for 00:05:00
9. Carefully remove supernatant.
10. Add 100 µL BAMBANKER **BioCat GmbH Catalog** #BB03-NP per 100,000 nuclei collected to the tube.
11. Gently resuspend.
12. Store in -80 °C freezer

#### *General Recommendations for the user*

3 For every new experiment we recommend performing the following steps:
When loading your tube into the FACS machine, run the unstained / IgG control sample first as this aids in setting the baseline parameters
Check your **event rate** in the **Acquisition Dashboard** window. If it is greater than 1500 evt/s turn down the "**flow rate**" or unload and dilute the sample further. If less than 100 evt/s, turn up the "**flow rate**" (don't exceed a flow rate of 5.0 if possible, as the instrument is less focused and more inaccurate at higher flow rates)
In the **Acquisition Dashboard** window choose the appropriate "**stopping gate**" and "**storage gate**" (when working with nuclei, set as "Nuclei" and "All events" respectively)
Choose the range of "**events to record**" and "**events to display**" that best suits your purpose (≥ 5000 for both is advised)
Under the **"threshold"** tab in the Cytometer window, change the threshold (should be set for FSC) so that any small events in the bottom corner of the FSC vs SSC graph (caused by general cell debris and dust) are no longer shown. The threshold should not be set too high so that it causes an arbitrary, artificial cut-off through the left side of your population but not so low that small events caused by debris/dust are visible (ideally between a threshold 200-500).
Under the **"parameters"** tab in the Cytometer window, adjust the "**FSC**" and "**SSC"** values to get your population sitting in the centre of the FSC vs SSC graph (a re-adjustment of the "**threshold**" may be required at this point). It is essential to select **"restart**" each time any of the parameters are changed to update the events being displayed to ensure only events are recorded under the new settings.
Adjust or draw a new gate in the FSC vs SSC plot to encompass the population of interest.
Look in the scatter graph of SSC-A vs SSC-W (if you opened a blank experiment you will need to draw one). Right-click on the graph and check it is only displaying the events encompassed by your previous FSC vs SSC gate. Adjust or draw a gate for SSC A vs SSC W to encompass all of the main population to the left of the graph and exclude outliers to the right (these are doublets and other cell debris clumps)
Under the **"parameters"** tab in the Cytometer window adjust parameters for the fluorochromes selected so the unstained / IgG control sample sits close to 0 for the fluorochrome on a graph of FSC vs fluorochrome.
Load the stained samples and check the stained population has a clear increase in signal for the fluorochrome in comparison to the unstained (signal should not exceed $10^{x4}$). Several minor re-adjustments of the fluorochrome's **"parameters"** may be necessary for the stained sample at this stage. If so, the unstained / IgG control has to be reset and re-recorded.

Note

**WARNING** – Do not change parameter settings between samples you wish to compare, if you do you will need to re-record all samples using the changed parameters.

11. Select the correct option for the collection device in the **Sort Layout** window ( we recommend "**4-Way Purity**" for general collection)
12. Regularly check your "**Efficiency**" in the **Sort Layout** window value. Between 80-100% is ideal, 70% is acceptable if less than 70% either the sample is too concentrated or you are sorting a rare population. Although the "**flow rate**" in the **Acquisition Dashboard** window can be increased to make the sort quicker, faster flow rates are less efficient.
13. Check the **"Electronic abort rate"** (N° errors /sec) and **"Electronic abort count"** (Tot N° of errors) at the bottom of the **Acquisition Dashboard** window. These parameters measure potential miss-sorts (different from efficiency as efficiency measures undetermined drops which are directed to the "**Waste**" and therefore lost but do not contaminate).**"Electronic abort rate"** should be <1% of total events per second.
14. For long sorts, gate positions should be regularly monitored, especially for stained populations as fluorochromes lose intensity over time and the population can shift towards the unstained. Gates can be moved during long sorts to compensate.
