## Supplementary material for "Optimised fluorescence-activated nuclei sorting for epigenomic analysis of cortical cell types": Protocol 2

Sep 29, 2023

### Purification of human cortex excitatory neuron nuclei from fetal and postnatal tissue using fluorescent activated nuclei sorting (FANS) in combination with a SATB2 antibody.

Forked from Fluorescence-activated nuclei sorting (FANS) on human post-mortem cortex tissue enabling the isolation of distinct neural cell populations for multiple omic profiling

DOI
dx.doi.org/10.17504/protocols.io.n92ldz9d8v5b/v1

Jonathan P Davies[1], Stefania S Policicchio[1], Barry Chioza[1], Gina Commin[1], Joe Burrage[1], Emma L Dempster[1], Jonathan Mill[1]

[1]University of Exeter Medical School, Exeter, UK

Complex Disease Epige...

Jonathan P Davies

OPEN ACCESS

**DOI:** https://dx.doi.org/10.17504/protocols.io.n92ldz9d8v5b/v1

**Protocol Citation:** Jonathan P Davies, Stefania S Policicchio, Barry Chioza, Gina Commin, Joe Burrage, Emma L Dempster, Jonathan Mill 2023. Purification of human cortex excitatory neuron nuclei from fetal and postnatal tissue using fluorescent activated nuclei sorting (FANS) in combination with a SATB2 antibody.. **protocols.io**
**https://dx.doi.org/10.17504/protocols.io.n92ldz9d8v5b/v1**

**License:** This is an open access protocol distributed under the terms of the **Creative Commons Attribution License**, which permits unrestricted use, distribution, and reproduction in any medium, provided the original author and source are credited

**Protocol status:** Working
**We use this protocol and it's working**

**Created:** January 27, 2022

**Last Modified:** September 29, 2023

**Protocol Integer ID:** 57511

**Keywords:** FANS, post-mortem brain, nuclei, flow cytometry, anti-NeuN, nuclei sorting, anti-SATB2, fetal, neurodevelopment, FACS, neuron, development, nuclei from multiple different human brain cell, optimal marker for neuronal nuclei, purification of human cortex excitatory neuron nuclei, neuronal nuclei, nuclei from excitatory neuron, cellular composition in regulatory genomic study, multiple different human brain cell, human cortex excitatory neuron nuclei, heterogeneous mix of different neural cell type, epigenetic process, satb2 antibody, fetal cortex, different neural cell type, postnatal cortex, regulatory genomic study, open chromatin analysis, gene regulation, current analyses of the human brain, genome, human brain, postnatal tissue, rna, activated nuclei, specific patterns of gene regulation, transcriptional variation in health, specific cell, purified populations of nuclei, transcriptional variation, gene expression, cellular composition, determining cell type, dna, excitatory neuron, neuron, nuclei, cortex, cell type

**Funders Acknowledgements:**
**Alzheimer's Research UK (ARUK)**
**Grant ID: ARUK-PPG2018A-010**
**Simons Foundation for Autism Research (SFARI)**
**Grant ID: 573312**
**Medical Research Council UK (MRC UK)**
**Grant ID: MR/R005176/1**

### Abstract

Increased understanding of the functional complexity of the genome has led to growing recognition about the role of epigenetic/transcriptional variation in health and disease. Current analyses of the human brain, however, are limited by the use of "bulk" tissue, comprising a heterogeneous mix of different neural cell types. Because epigenetic processes play a critical role in determining cell type-specific patterns of gene regulation it is important to consider cellular composition in regulatory genomic studies of human post-mortem tissue, and there is a need for methods to purify populations of specific cell-types. This protocol builds on a **previous protocol** that uses fluorescence-activated nuclei sorting (FANS) to isolate and profile nuclei from multiple different human brain cell-types from frozen post-mortem tissue. Because NeuN is not an optimal marker for neuronal nuclei from fetal cortex, we have optimized a method using a SATB2 antibody to purify nuclei from excitatory neurons in both fetal and postnatal cortex. Purified populations of nuclei are amenable to simultaneous profiling of i) DNA modifications (via bisulfite sequencing / array), ii) histone modifications (via CUT&Tag), iii) open chromatin analysis (via ATAC-seq), and iv) gene expression (via RNA-seq).

| | | |
|---|---|---|
| | **Supplier** | Thermo Scientific™ |
| | **Model** | Sorvall™ WX 80+ |
| | **Rotor** | TH-641 |
| | **Speed** | 25,200 RPM / 108670.8 x g |
| | **Acceleration** | 9 |
| | **Deceleration** | 5 |
| | **Temperature** | 4°C |

***Table 4:*** *Ultracentrifuge specification and conditions*

| | A | B | C | D | E |
|---|---|---|---|---|---|
| | **Antibody** | **Preconjugated** | **Supplier** | **Cat No** | **Dilution** |
| | Hoechst 33342 | -- | Abcam | ab228551 | 1:500 |
| | Anti-SATB2 | to Alexa Fluor 488 | Abcam | ab196316 | 1:1000 |
| | Anti-NeuN | to Alexa Fluor488 | Millipore | MAB377X | 1:1000 |

***Table 5***: List of antibodies required for FANS protocol

#### Troubleshooting

#### Nuclear prep for FACS separation (using SATB2 and Hoechst)

1 The protocol below yields approximately 660,000 SATB2$^{+ve}$, 800,000 SATB2 $^{-ve}$ nuclei per 200 mg of frozen human fetal post-mortem cortex tissue. Recovery might vary from sample to sample due to high inter-sample variability (developmental stage and density brain collection, cortex sub-areas, fat content of tissue sectioned, and white to grey matter ratio). Yield will also vary for NeuN against developmental age, and we have shown NeuN is not suitable in pre-natal samples, but can be used in post-natal and adult cortex to collect a comparable population.

Refer to Materials-***Table 1*** for details about the equipment required and to Materials-***Table 2*** for specifications of reagents required.

Note

Throughout this protocol, SATB2 and NeuN are interchangeable as the neuronal antibody marker, depending on age of the individual being profiled.

1.1 ***Solution and buffer preps***

- *Lysis Buffer (LB)*
- *Sucrose Solution (SS)*
- *Staining Buffer (SB)*

Solutions should be kept at 4 °C or On ice. Refer to Materials-***Table 3*** for recipes of solutions and buffers.

Note

LB and SS can be prepared a week in advance, with DTT added on the day of use. Solutions should be stored at 4 °C once made, preferably in brown glass bottles.

Note

SB should be prepared fresh each day.

Note

Samples are homogenised as bulk tissue using a 7 mL Dounce homogeniser and then equally divided into two ultracentrifuge tubes.

1.2 ***Nuclei isolation*** 1h 20m

1. Pre-cool the ultracentrifuge, including the rotor and swing buckets, to 4 °C 00:30:00 before starting this stage of the protocol.
2. All buffers and the Dounce homogenisers should be pre-cooled on ice.
4. Add DTT ( [M] 1 millimolar (mM) final concentration ) to the SS and LB (i.e. 17 µL 3M DTT per 50 mL of SS/LB)
5. Transfer 2 mL LB to the homogeniser per 200 mg human brain tissue
6. Add the dissected tissue sample into the homogeniser
7. Wait 00:05:00 before douncing the tissue to allow the sample to defrost

To reduce heat caused by friction, the Dounce homogenisation step should be performed on ice with gentle strokes, and care should be taken to avoid foaming.

Note

Using the "TIGHT" pestle helps reduce the number of strokes required to reach full tissue disruption.

Note

The number of strokes required to fully homogenise the tissue may vary between samples due to heterogeneity in cellular composition, lipid content, and the amount of connective tissue.

Note

In comparison to adult tissue, the fetal tissue dissociates very easily, with fewer strokes required, and the lysate will be very pale. However, the cells are densely packed in a fetal brain, so we retained the same volume of lysis buffer to perform the dissociation and staining protocols.
Because the fetal brain is more easily dissociated, fewer strokes are required, and we have even seen success with gentle pipetting to agitate the tissue, rather than using a dounce.

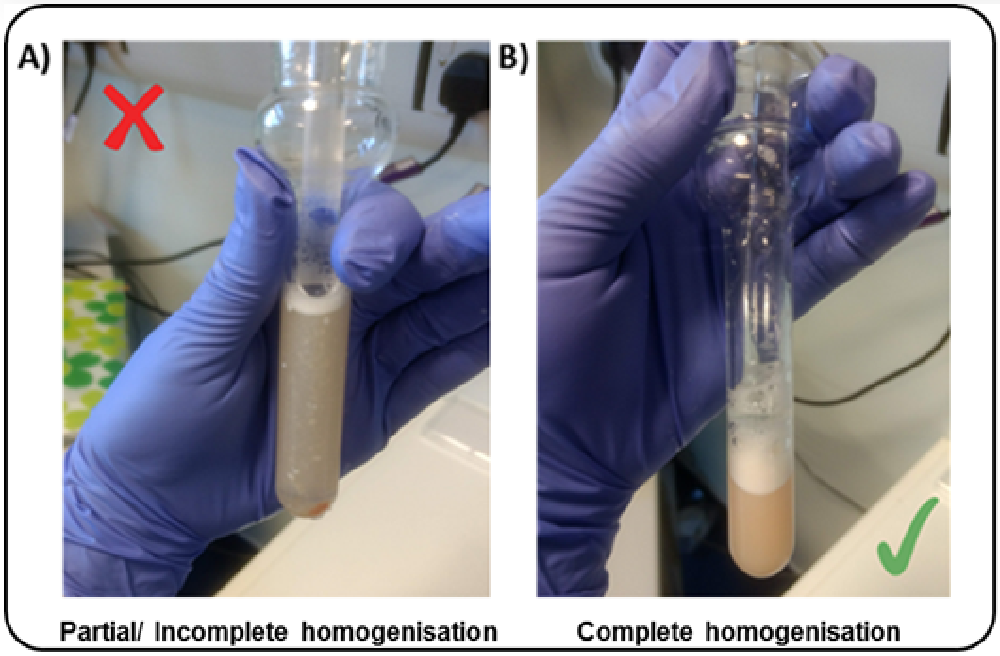

***Figure 1*** Example of adult brain tissue sample
**A)** only partially homogenised **B)** complete homogenisation.
Fetal samples will appear paler, with fewer pieces of floating lipid based tissue.

8.Transfer 8 mL SS (1.8M) to PA thin-walled ultracentrifuge tubes

9.Carefully overlay with tissue homogenate (1 mL per tube) - using a P1000 pipette, releasing slowly down the side of the tube

10.Overlay with another 1 mL LB – do not worry about disrupting the homogenate phase
11.Balance opposite tubes by weight with 1x PBS using a fine microbalance
12.Perform ultracentrifugation for 00:45:00 (see ***Table 4*** for centrifuge specification and conditions)

1.3 ***After Ultracentrifugation step***

1. Pour off any the supernatant, taking care not to dislodge the pellet (90-degree inclination of the tube). If the pellet is hard to see, it is okay to leave 100-200 µL solution in the ultracentrifuge tube
2. Re-suspend pellet in SB ( 750 µL ), gently pipette up and down
3. Let samples sit on ice for 00:15:00 at least ( **Blocking step** )
4. Transfer volume into 2 mL tubes
5. Rinse out ultracentrifuge tubes in order to maximise nuclei collection by adding 750 µL of SB per tube, pipetting up and down several times, and transferring into the 2 mL tubes
6. **Centrifuge step:** for 00:05:00 , 4 °C
7. Discard supernatant (pipetting off, or pouring off gently)
8. Re-suspend each nuclei pellet in fresh SB (500 µL)
9. If the sample was split then pool together pellets from the same sample (Final Volume = 1 mL )
10. Add DNA dye (Hoechst, 1:1000) and mix thoroughly via inversion.
11. Pipette out 100 µL of nuclei solution for the Unstained Control (Hoechst dye only) and transfer to a new 2 mL tube
12. Bring the volume up to

500 µL
for the Unstained tube with fresh SB
13. Replace the 100µL taken from the Stained" tube with 100 µL of fresh SB (Final Volume =
1 mL 1 mL )

Note

We don't add 1 mL of SB to the ultracentrifuge pellet, as there is always residual liquid remaining, and this would push the volumes above the 2 mL maximum in the 2 mL tubes.

Note

The amount of sample removed for the unstained control can vary, if you're concerned about wasting material. It is only used to establish the gating thresholds, or to collect a non-sorted population.

1.4 ***Immunostaining***

1. Add your antibody of choice (Ab) to staining tube:

SATB2 Alexa488 (1:1000)

or

NeuN Alexa488 (1:1000)

Refer to ***Table 5*** for specifications of the antibodies used

2. Incubate tubes for 01:30:00 on the rotor (speed=14 max) at 4 °C , keeping the tubes in the dark
3. Washing step: 1000 x g for 00:05:00 , 4 °C (both "Stained" and "Unstained" tubes)
4. Discard supernatant (by pipetting off)
5. Re-suspend in fresh SB ( 1 mL for the Unstained tube, 1.5-2 mL for the Stained tube - depending on pellet size)

#### Fluorescence-Activated Nuclei Sorting (FANS)

5m

2 For machine start-up, CST and Accudrop calibrations refer to **BD FACSAria III User's Guide** for guidance and troubleshooting. The following instructions describe FANS using BD FACSAria III. Other FACS platforms can be used but might require modifications to the protocol.

2.1 ***General Gating Parameters***

For each sample, load stained and unstained tubes individually for data acquisition. A preliminary qualitative analysis of the acquired data is essential to select the appropriate gating strategy to maximize the nuclei capture while excluding unnecessary debris and to ensure optimal signal/noise ratio.

*Gating Parameters (X-axis:Y-axis):*

FSC-A:SSC-A (Size, cell granularity or internal complexity)

SSC-W:SSC-A (to gate out doublets)

FSC-A:DAPI-A (to gate the single nuclei population)

DAPI-A:FITC-A (to gate SATB2 or NeuN stained nuclei)

Note

Refer to **Figure 2** for a visualization of the gating strategy. Gating is conserved as much as possible across the age trajectory, but can vary due to developmental changes. It's important to note that the populations should still be present when using SATB2

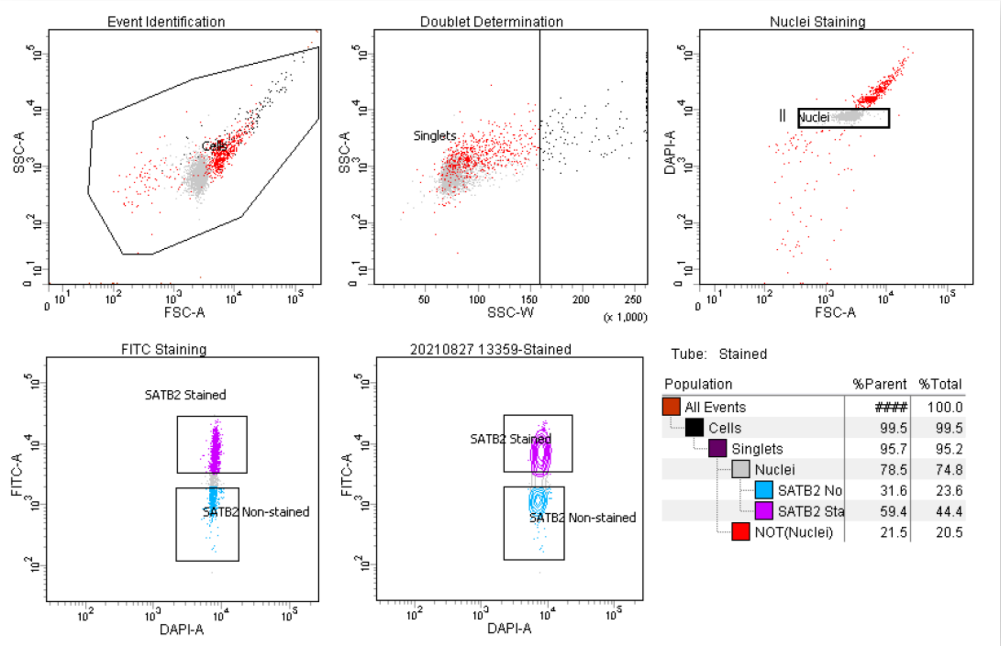

**Figure 2.** FANS gating strategy on a fetal sample 13 post conception weeks of development (13 pcw).
**(a)** Particles smaller than nuclei (black dots) were eliminated with an area plot of forward-scatter (FSC-A) versus side-scatter (SSC-A), with gating for nuclei-sized particles inside the gate (box).
**(b)** Plots of height versus width in the side scatter channel are used for doublet discrimination with gating to exclude aggregates of two or more nuclei.
**(c)** Doublet discrimination gating was used to isolate nuclei determined by subgating on Hoechst 33342. (d, e,) Subsequent scatterplots discerning
**(d)** SATB2-Alexa Fluor488–conjugated antibody staining (purple) for SATB2 stained neuronal cells and SATB2 non-stained nuclei (blue) for non-neuronal cells
**(e)** SATB2-Alexa Fluor488–conjugated antibody staining (purple) in a contour plot.

#### 2.2 ***Data recording settings***

In line with the experiment design, FSC, SSC, DAPI, FITC, are the parameters for which voltage values may need to be slightly adjusted due to experiment/ inter-sample variability.

It is advisable to set the threshold value at 200 during data recording. Moreover, in the acquisition dashboard tab, we recommend setting **Events to Record** ≤3000, **Event to display** ≤1000 and **Flow Rate** = 1.0 (1,000 events per second) in order to increase the accuracy of signal detection.

The flow rate can be increased during sample collection to reduce the sort speed (ideally max events per second =1,500 for a 100-micron nozzle). However higher flow rates impact the data resolution and accuracy of events detection, and subsequent sorting of cellular fractions (see **BD FACSAria III User's Guide** for details).

During analysis, recorded data is displayed in plots, while gates are used to define populations of interest for selection. **Figure 3** shows a representative example of the two most common outcomes we often observe.

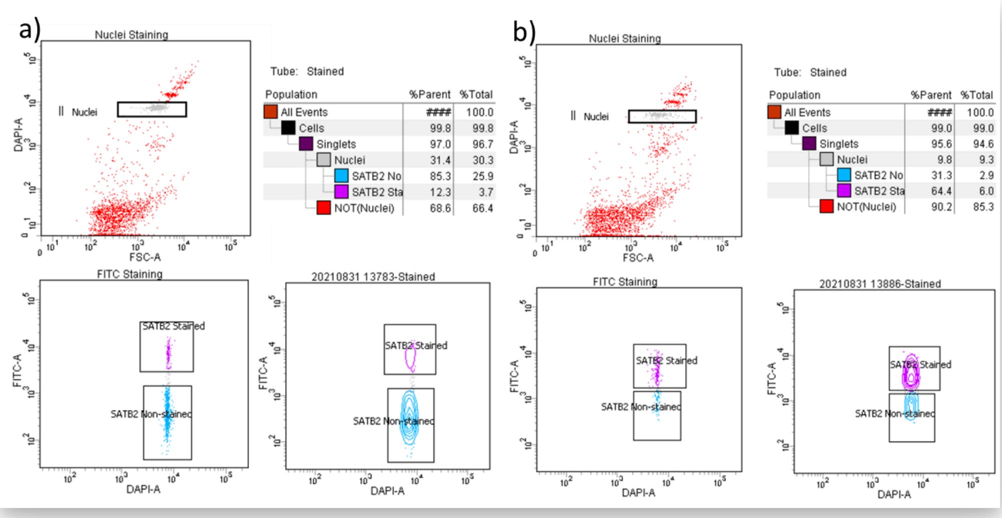

***Figure 3**. Representative example of inter-individual variability.*

The data shown here are derived from two different control prefrontal cortex specimens of comparable age, gender, and brain collection which were processed in parallel following the same procedure.

**a)** sample from 13pcw, clean nuclei isolation, but with smaller SATB2$^{+ve}$ fraction with clear separation of populations

**b)** poor sample separation on a 20pcw sample, but contour map demonstrates distinct populations.

2.3 ***Sample Collection*** 5m

1. 1. LoBind Tubes (Eppendorf, Cat No:30108051) are required to collect nuclei (to maximize sample recovery of nucleic acids by significantly reducing sample–to–surface binding).
2. 2. Collected fractions can be used directly for downstream applications (e.g. DNA/RNA extraction, chromatin shearing).
3. 3. If you are collecting for DNA or RNA, as soon as the number of events desired is reached, transfer the tubes On ice , do not hold them at Room temperature .

Note

During collection, it is crucial to regularly pause the sorting to mix the two phases in order to preserve the integrity of resulting RNA preparations.

Note

For RNA extraction, LoBind Tubes should each contain 1 mL of pre-chilled TRIzol™ LS Reagent (Fisher Scientific, Cat No: 11578616) prior to sorting. If possible, use a chilled collection tube holder on your FACS machine.

4. Keep samples On ice and in the dark for the entire duration of the sorting.
5. Lightly vortex sample tubes to make the mixture homogeneous (not clumped) before loading the tube into the FACS chamber.
6. Load the **UNSTAINED** control tube into the chamber first to postion Unstained gating boxes.
7. Proceed by collecting **STAINED** tube by simultaneously sorting for SATB2$^{+ve}$ (positive) or SATB2$^{-ve}$ (negative) and proceed with nuclei collection.

Note

We observe great variability of staining profile in the fetal samples across the trajectory of development. We advise you to not expect as conserved patterns as observed in adult cortex.

8. For long term storage of collected nuclei 1000 x g, 4°C for 00:05:00
9. Carefully remove supernatant.
10. Add 100 µL BAMBANKER to the tube.
11. Gently resuspend.
12. Store in -80 °C freezer
