## Supplementary material for "Optimised fluorescence-activated nuclei sorting for epigenomic analysis of cortical cell types": Protocol 3

Sep 04, 2025 Version 2

### Fluorescence-activated nuclei sorting (FANS) of purified neural cell populations from mouse cortex for multi-omic profiling V.2

Version 1 is forked from Fluorescence-activated nuclei sorting (FANS) on human post-mortem cortex tissue enabling the isolation of distinct neural cell populations for multiple omic profiling

DOI
dx.doi.org/10.17504/protocols.io.dm6gpbwndlzp/v2

Stefania S Policicchio[1], Isabel Castanho[1], Barry Chioza[1], Joe Burrage[1], Jonathan Mill[1], Emma L Dempster[1], Jonathan P Davies[1]

[1]University of Exeter Medical School, Exeter, UK

Complex Disease Epige...

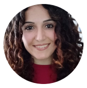

Stefania S Policicchio
University of Exeter Medical School

OPEN ACCESS

**DOI:** https://dx.doi.org/10.17504/protocols.io.dm6gpbwndlzp/v2

**Protocol Citation:** Stefania S Policicchio, Isabel Castanho, Barry Chioza, Joe Burrage, Jonathan Mill, Emma L Dempster, Jonathan P Davies 2025. Fluorescence-activated nuclei sorting (FANS) of purified neural cell populations from mouse cortex for multi-omic profiling. **protocols.io** **https://dx.doi.org/10.17504/protocols.io.dm6gpbwndlzp/v2**Version created by **Barry Chioza**

**License:** This is an open access protocol distributed under the terms of the **Creative Commons Attribution License**, which permits unrestricted use, distribution, and reproduction in any medium, provided the original author and source are credited

**Protocol status:** Working
**We use this protocol and it's working**

**Created:** August 14, 2025

**Last Modified:** September 04, 2025

**Protocol Integer ID:** 224702

**Keywords:** FANS, nuclei, flow cytometry, anti-NeuN, nuclei sorting, mouse cortex, anti- PU.1, nuclei from multiple different cell type, cellular composition in regulatory genomic study, purified neural cell population, other glial origin nuclei, frozen mouse cortex tissue, neural cell populations from mouse cortex, chromatin accessibility, epigenetic process, regulatory genomic study, purified populations of nuclei, heterogeneous tissue like the brain, different cortical cell type, microglia, frozen mouse cortex, specific patterns of gene regulation, gene regulation, multiple different cell type, genome, mouse cortex, histone modification, transcriptional variation in health, microarray, transcriptional variation, activated nuclei, determining cell type, dna modification, rna, populations of nuclei, dna, gene expression, cell type, cellular composition

#### Abstract

Increased understanding of the functional complexity of the genome has led to growing recognition about the role of epigenetic/transcriptional variation in health and disease. Because epigenetic processes play a critical role in determining cell type-specific patterns of gene regulation it is important to consider cellular composition in regulatory genomic studies of heterogeneous tissue like the brain. Building on a **previous protocol** for isolating purified populations of nuclei from different cortical cell types from human post-mortem brain tissue, this protocol uses fluorescence-activated nuclei sorting (FANS) to isolate and profile nuclei from multiple different cell types from frozen mouse cortex. This protocol can be used to robustly purify populations of neuronal (NeuN$^{+ve}$) and microglia (PU.1$^{+ve}$) and other glial origin nuclei (NeuN$^{-ve}$/PU.1$^{-ve}$) from frozen mouse cortex tissue, with each sample yielding purified populations of nuclei amenable to simultaneous analysis of i) DNA modifications (via bisulfite sequencing/microarray), ii) histone modifications, iii) chromatin accessibility (via ATAC-seq), and iv) gene expression (via RNA-seq).

#### Materials

| | A | B | C |
|---|---|---|---|
| | **Equipment** | **Supplier** | **Catalogue No** |
| | **BD FACSAria™ III Cell Sorter** | BD Biosciences | 648282-23 |
| | **Sorvall WX 80+ Ultracentrifuge** | Thermo Scientific™ | 75000080 |
| | **1mL Dounce Tissue Grinder** | Sigma-Aldrich | DWK885300-0001-1EA |
| | **PA Thin-walled ultracentrifuge tubes** | Thermo Scientific | 03699 |

| | A | B | C |
|---|---|---|---|
| | **5% BSA Solution (BB)** | | |
| | BSA | | 200 mg |
| | 1x PBS | | 4 mL |
| | **Staining Buffer (SB)** | | |
| | 0.5% BSA | 5% BSA Solution (BB) | 400 µL |
| | 10X PBS | | 400 µL |
| | *Optional: RiboLock RNase Inhibitor 0.2U/ µL* | 40U/ µL | 5 µL / 1 mL |
| | Adjust with ddH2O to | | 4 mL |

**Table 3** : Recipes for buffers and solutions required

| | A | B |
|---|---|---|
| | **Supplier** | Thermo Scientific‰ |
| | **Model** | Sorvall‰ WX 80+ |
| | **Rotor** | TH-641 |
| | **Speed** | 25,200 RPM / 108670.8 x g |
| | **Acceleration** | 9 |
| | **Deceleration** | 5 |
| | **Temperature** | 4°C |

***Table 4*** *:* Ultracentrifuge specification and conditions

### Troubleshooting

#### Nuclear prep for FACS separation (using NeuN, PU.1 and Hoechst)

2h 40m

1 In our hands the protocol below yields ~60,000 $NeuN^{+ve}$ (neuron enriched), ~5,000 $PU.1^{+ve}$ (microglial enriched) and ~20,000 double negative ($NeuN^{-ve}$/$PU.1^{-ve}$; oligodendrocyte enriched) nuclei per ≤ 100 mg of frozen mouse cortex tissue. Recovery might vary from sample to sample due to high inter-sample variability and regional differences across cortical areas (i.e. dissection procedure, fat content of tissue sectioned, age).

Refer to **Materials-Table 1** for details about the equipment required and to **Materials-Table 2** for specifications of reagents required.

1.1 ***Solution and buffer preps***

- *Lysis Buffer (LB)*
- *Sucrose Solution (SS)*
- *Staining Buffer (SB)*

Solutions should be kept at 4 °C or On ice . Refer to **Materials -Table 3** for recipes of solutions and buffers.

Note

**NOTE 1** – LB and SS can be prepared a week in advance, with DTT added on the day of use. Solutions should be stored at 4 °C once made.

Note

**NOTE 2** – SB should be prepared fresh each day.

Note

**NOTE 3**- Samples are homogenised as bulk tissue using a 1 mL Dounce homogeniser and then equally divided into two ultracentrifuge tubes.

1.2 ***Nuclei isolation***

45m

1.Pre-cool the ultracentrifuge to 4 °C before starting this stage of the protocol.

2.All buffers and the Dounce homogenisers should be pre-cooled on ice.

4. Add 1 mM DTT to the SS and LB according to the recipe (i.e. 17 µL 3M DTT per 50 mL of SS/LB).

5. Transfer 1 mL LB into each homogeniser.

6. Add the dissected tissue sample into the homogeniser.
7. Wait 3-5 minutes before douncing the tissue to allow the sample to defrost.
To reduce heat caused by friction, the Dounce homogenisation step should be performed on ice with gentle strokes, and care should be taken to avoid foaming.

Note

**NOTE 1** – Using the "TIGHT" pestle helps reduce the number of strokes required to reach full tissue disruption.

Note

**NOTE 2** - The number of strokes required to fully homogenise the tissue may vary between samples due to heterogeneity in cellular composition, lipid content, and the amount of connective tissue.

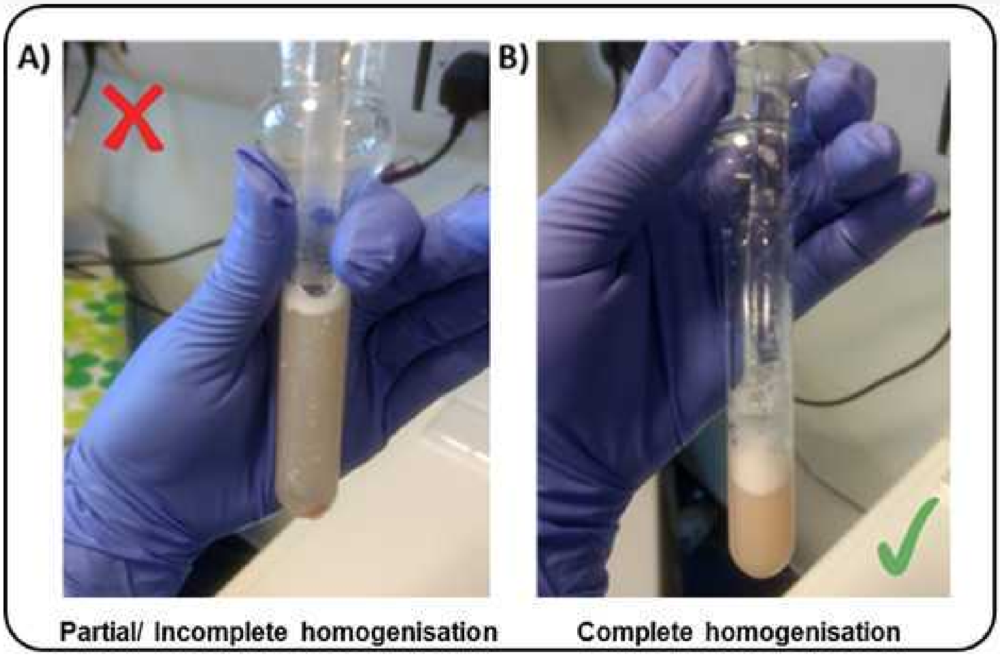

***Figure 1*** Example of brain tissue sample A) only partially homogenised B) complete homogenisation

8. Transfer 8 mL SS (1.8M) to PA thin-walled ultracentrifuge tubes.
9. Carefully overlay with tissue homogenate (1 mL per tube) - using a P1000 pipette, releasing slowly down the side of the tube.
10. Add 1mL LB back into the homogeniser to rinse and recover as much as possible of the residual homogenate left behind.
11. Recover volume from homogeniser and overlay it on homogenate layer.
12. Balance opposite tubes by weight with 1x PBS using a fine microbalance.

13. Perform ultracentrifugation for 00:45:00 (see **Table 4** for centrifuge specification and conditions).

1.3 ***After Ultracentrifugation step*** 20m

1. Aspirate supernatant leaving 1-2 mL of the solution in the tube along with the pellet.
2. Pour off any remaining supernatant, taking care not to dislodge the pellet (90-degree inclination of the tube). If the pellet is hard to see, it is okay to leave 100-200 µL solution in the ultracentrifuge tube
3. Resuspend pellet in SB (1 mL), gently pipette up and down.
4. Let samples sit On ice for 00:15:00 at least **(Blocking step).**
5. Transfer volume into a new 1.5 mL LoBind DNA tube.
6. Rinse out ultracentrifuge tubes in order to maximise nuclei collection by adding 1 extra mL of SB per tube, pipetting up and down several times, and transferring into a new 1.5 mL LoBind DNA tube.
7. **Washing step:** 1000 x g, 4°C for 00:05:00
8. Discard supernatant (pipetting off gently).
9. Re-suspend each nuclei pellet in fresh SB (500 µL).
10. If the sample was split then pool together resuspended pellets from the same sample (Final Volume = 1 mL)
11. Add DNA dye (Hoechst, 2 µL/1 mL) and mix thoroughly by inversion.
12. Pipette out 120 µL of nuclei solution for the Unstained Control (Hoechst dye only) and transfer to a new 1.5 mL LoBind DNA tube.
13. Bring the volume up to 500 µL for the Unstained tube with fresh SB.
14. Replace the volume taken from the "Stained" tube with 120 µL of fresh SB (Final Volume = 1 mL).

1.4 ***Immunostaining*** 1h 35m

1. Add the following three antibodies (Ab) to the "Stained" tube (1mL final volume):
- PE anti-PU.1 (1:100 dilution) – [10 µL Ab]
- Alexa488 anti-NeuN (1:1000 dilution) – [1 µL Ab ]

Refer to **Table 5** for specifications of the antibodies used

2. Incubate tubes for 01:30:00 on the rotor (speed=11) at 4 °C , keeping the tubes in the dark.
3. Washing step: 1000 x g, 4°C for 00:05:00 (both "Stained" and "Unstained" tubes).
4. Discard supernatant (by pipetting off).
5. Re-suspend in fresh SB (500 µL for the Unstained tube, 1 mL for the Stained tube - depending on pellet size).

#### Fluorescence-Activated Nuclei Sorting (FANS)

2 For machine start-up, CST and Accudrop calibrations refer to **BD FACSAria III User's Guide** for guidance and troubleshooting. The following instructions describe FANS using BD FACSAria III. Other FACS platforms can be used but might require modifications to the protocol.

2.1 ***General Gating Parameters***

For each sample, load stained and unstained tubes individually for data acquisition. A preliminary qualitative analysis of the data acquired is essential to select the appropriate gating strategy to maximise the nuclei capture while excluding unnecessary debris and to ensure an optimal signal/noise ratio.

*Gating Parameters (X-axis: Y-axis):*

**FSC-A:SSC-A** (Size, cell granularity or internal complexity)
**SSC-W:SSC-A** (to gate out doublets)
**FSC-A:DAPI-A** (to gate the single nuclei population)
**DAPI-A:FITC-A** (to gate NeuN stained nuclei)
**DAPI-A:PE-A** (to gate PU.1 stained nuclei)
**FITC-A: PE-A** (to visualize the distribution of the three detectable populations)

Note

The PU.1+ve population (microglia) is gated as a "daughter" population from the NeuN-ve fraction (non neuronal nuclei). Refer to **Figure 2** for a visualization of the gating strategy.

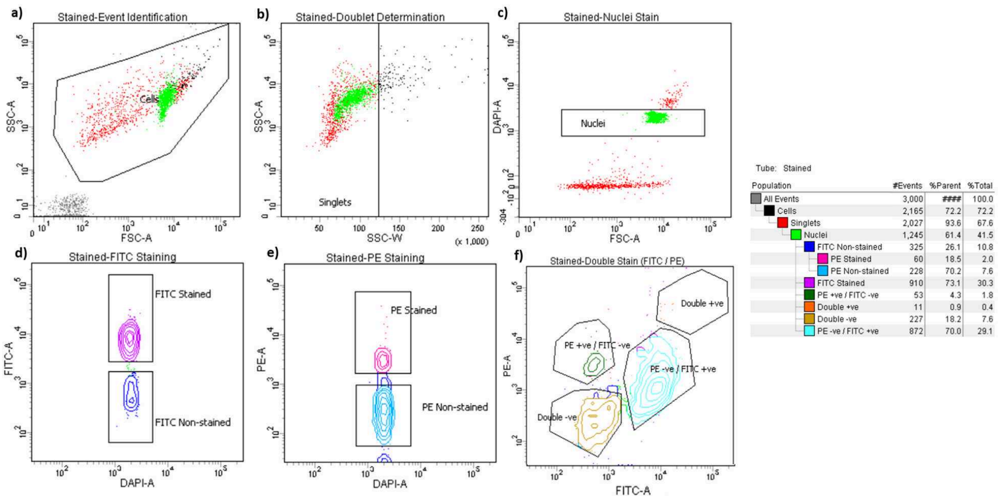

***Figure 2.*** FANS gating strategy. **(a)** Particles smaller than nuclei (black dots) were eliminated with an area plot of forward-scatter (FSC-A) versus side-scatter (SSC-A), with gating for nuclei-sized particles inside the gate (box). **(b)** Plots of height versus width in the side scatter channel are used for doublet discrimination with gating to exclude aggregates of two or more nuclei. **(c)** Doublet discrimination gating was used to isolate nuclei determined by sub-gating on Hoechst 33342. (d-f) Subsequent scatterplots discerning **(d)** NeuN-Alexa Fluor488–conjugated antibody staining (purple) **(e)** PU.1 PE-stained nuclei (dark pink) **(f)** the distribution of the three main nuclei subpopulations identified through double staining strategy (NeuN$^{+ve}$, neurons; PU.1$^{+ve}$, microglia, double$^{-ve}$, oligodendrocytes enriched). The resultant hierarchical colour key ensures that only nuclei that are negative for staining with the NeuN antibody are passed through the next gating condition.

2.2 ***Data recording settings***

In line with the experiment design, FSC, SSC, DAPI, FITC and PE are the parameters for which voltage values may need to be slightly adjusted due to experiment/ inter-sample variability.

It is advisable to set the threshold value between 200 and 500 during data recording. Moreover, in the acquisition dashboard tab, we recommend setting **Events to Record**≤3,000, **Event to display**≤5000 and **Flow Rate** = 1.0 (1,000 events per second) in order to increase the accuracy of signal detection.

The flow rate can be increased during sample collection to reduce the sort speed (ideally max events per second =1,500 for a 100-micron nozzle). However higher flow rates impact the data resolution and accuracy of events detection, and subsequent sorting of cellular fractions (see BD FACSAria III User's Guide for details).

During analysis, recorded data is displayed in plots, while gates are used to define populations of interest for selection. **Figure 3** shows a representative example of the two most common outcomes we often observe.

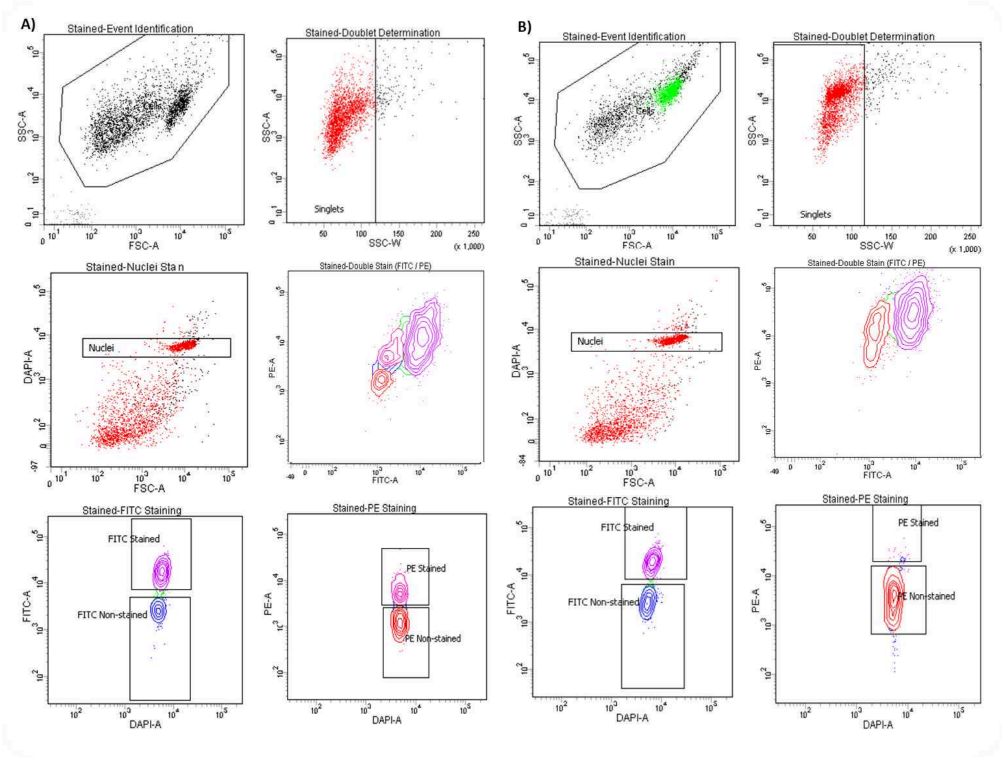

***Figure 3***. Representative example of inter-individual variability. The data shown here are derived from two different cortex specimens of comparable age, sex, and group which were processed in parallel following the same procedure. **A)** distinctly separated PU.1 +ve fraction vs **B)** missing PU.1+ve positively stained population.

2.3 **Sample Collection**

1. LoBind Tubes (Eppendorf, Cat No:30108051) are required to collect nuclei (to maximise sample recovery of nucleic acids by significantly reducing sample–to–surface binding).
2. Nuclei are collected in 150 µL BAMBANKER (up to 1,000,000 nuclei) if collecting for chromatin or DNA assays. Nuclei are collected in Trizol LS Reagent (500 µL) if you are collecting for RNA.
3. Collected fractions can be used directly for downstream applications (e.g. DNA/RNA extraction, chromatin shearing) or frozen at -80 °C for long term storage. We recommend not to pellet nuclei down prior freezing to prevent nuclei loss due to nuclei bursting/ damage.

Note

**NOTE -** During collection, it is crucial to regularly pause the sorting to mix the two phases by inversions in order to preserve the integrity of resulting RNA preparations.
For RNA extraction, LoBind Tubes should each contain 500µL of pre-chilled TRIzol™ LS Reagent (Fisher Scientific, Cat No: 11578616) prior to sorting.

4. Keep samples on ice for the entire duration of the sorting
5. Lightly vortex sample tubes to make the mixture homogeneous (not clumped) before loading the tube into the FACS chamber
6. Load the UNSTAINED control tube into the chamber first and proceed with nuclei collection (for DNA, 200,000 events; for RNA, 300,000 events; for ATAC-seq 50,000 events).
7. Proceed by collecting STAINED tube by simultaneously sorting for NeuN and PU.1
