## Supplementary Figures for "Optimised fluorescence-activated nuclei sorting for epigenomic analysis of cortical cell types"

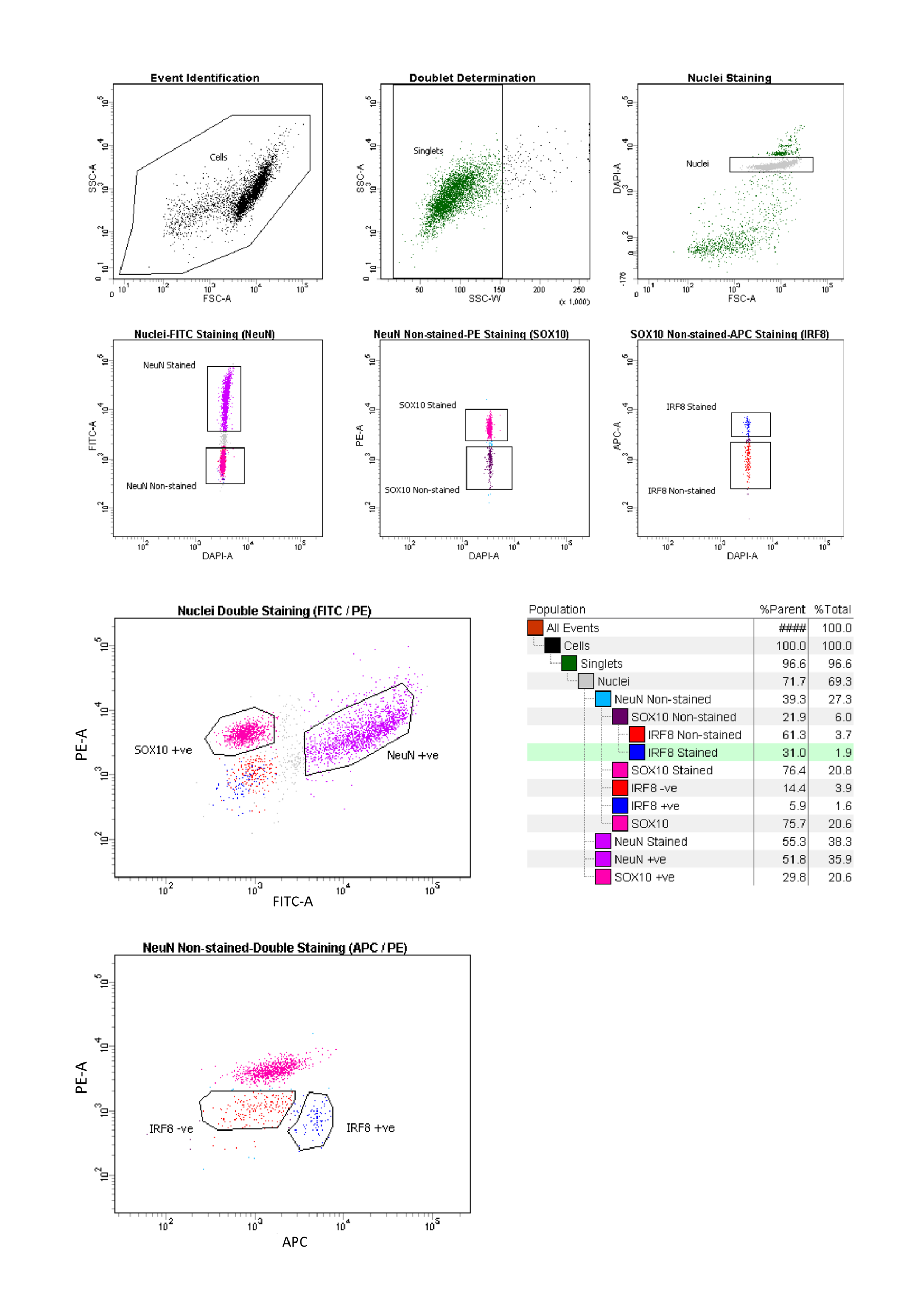

**(a)**

**(b)**

**(c)**

**(d)**

**(e)**

**(f)**

**(g)**

**(h)**

i

**(i)**

**Figure S1. FANS gating strategy for post-mortem postnatal human cortex.**

(**a**) Nuclei were first identified based on forward scatter (FSC-A) versus side scatter (SSC-A), with gates drawn to capture nuclei-sized particles. (**b**) Doublet discrimination was performed using side-scatter height versus width (SSC-H vs SSC-W), allowing exclusion of aggregates and selection of singlets. (**c**) Hoechst 33342 fluorescence was used to further refine the singlet nuclei population. **(d–f**) Sequential gating was then applied to identify nuclei positive for each antibody: (**d**) NeuN–Alexa Fluor 488 (neurons), (**e**) SOX10–NL577 (oligodendrocytes), and (**f**) IRF8–APC (microglia). (**g-h**) A combined triple-staining gate was used to classify nuclei into the major cortical populations: NeuN⁺ neurons, SOX10⁺ oligodendrocytes, IRF8⁺ microglia, and triple-negative (NeuN⁻/SOX10⁻/IRF8⁻) astrocyte-enriched nuclei. (**i**) The hierarchical gating strategy ensures that nuclei are assigned to mutually exclusive populations based on their NeuN, SOX10, and IRF8 staining status, with proportions calculated relative to the total nuclei gate.

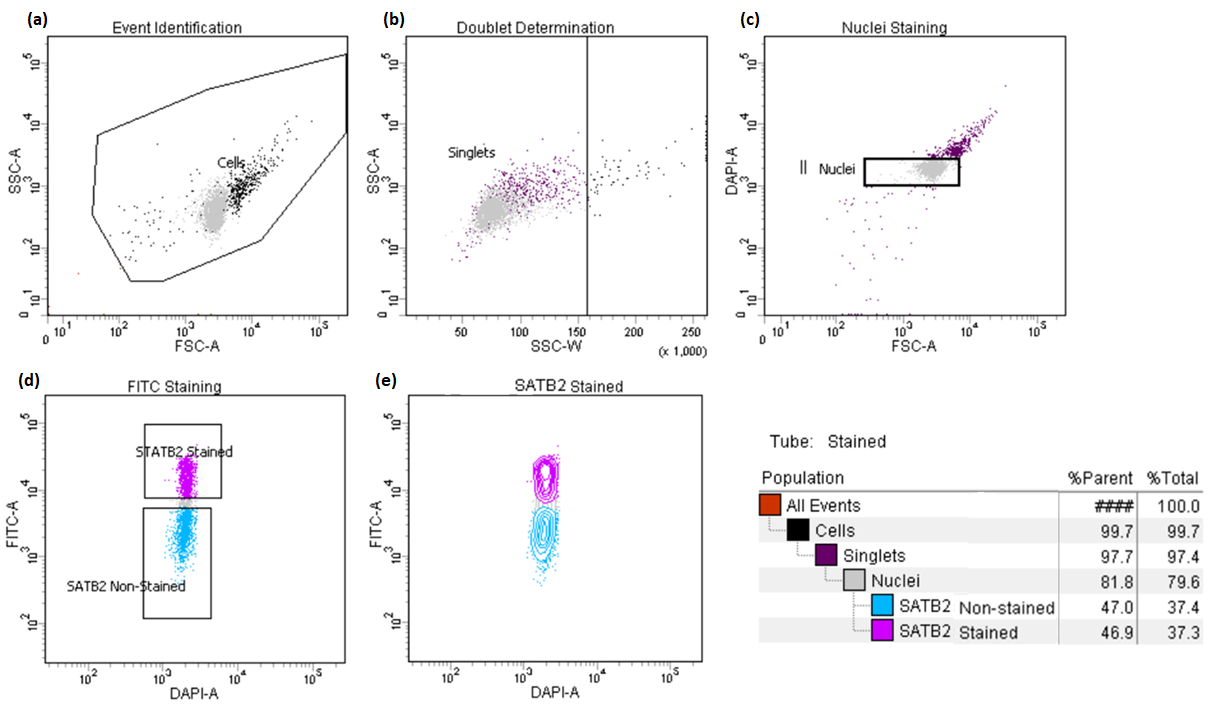

**Figure S2. Representative FANS gating strategy for post-mortem fetal cortex**.

(**a**) Nuclei were first distinguished from smaller debris by gating nuclei-sized particles in a forward scatter (FSC-A) versus side scatter (SSC-A) plot. (**b**) Side-scatter height versus width (SSC-H vs SSC-W) was used for doublet discrimination, allowing exclusion of aggregates. (**c**) Hoechst 33342 fluorescence was then used to refine the singlet nuclei population. (**d–e**) SATB2–Alexa Fluor 488 staining was used to identify excitatory neuron–enriched nuclei, shown in both (d) scatter and (e) contour plots. SATB2⁺ nuclei (purple) were clearly separated from the SATB2⁻ population (dark blue). The hierarchical colour key illustrates the gating workflow and reports the proportion of events assigned to each gate.

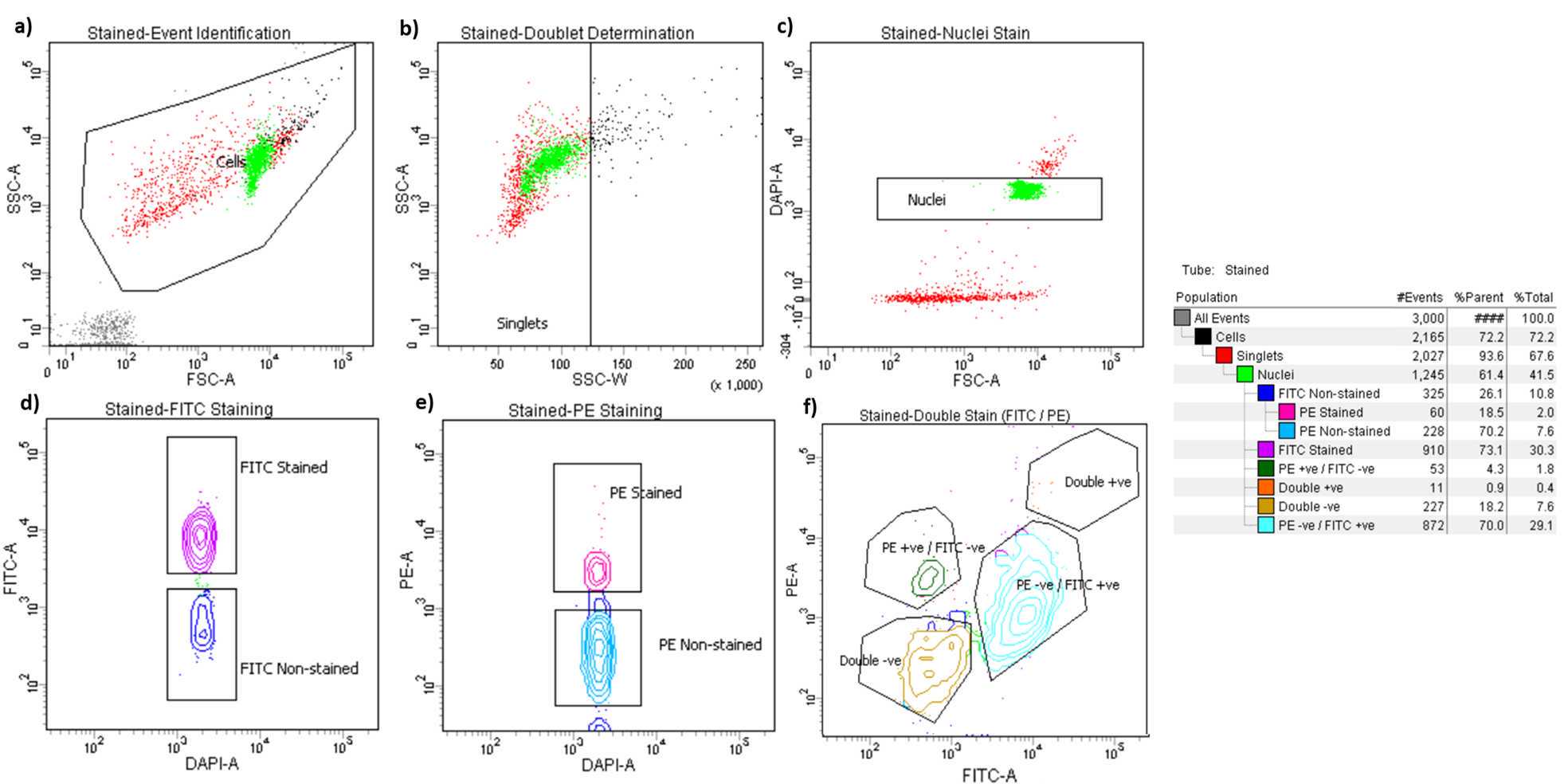

**Figure S3. Representative FANS gating strategy for post-mortem mouse cortex.**

(**a**) Nuclei were identified by gating nuclei-sized particles in a forward scatter (FSC-A) versus side scatter (SSC-A) plot, excluding smaller debris. (**b**) Side-scatter height versus width (SSC-H vs SSC-W) was used for doublet discrimination to remove aggregates. (**c**) Hoechst 33342 fluorescence was applied to further refine the singlet nuclei population. (**d-f**) Subsequent scatterplots discerning **(d)** NeuN-Alexa Fluor488–conjugated antibody staining (purple) **(e)** PU.1 PE-stained nuclei (dark pink) **(f)** the distribution of the three main nuclei subpopulations identified through double staining strategy (NeuN^+^, neurons; PU.1^+^, microglia, double^–^, oligodendrocytes enriched). The resultant hierarchical colour key ensures that only nuclei that are negative for staining with the NeuN antibody are passed through the next gating condition.

**Figure S4. Immunofluorescence imaging of NeuN- and SOX10-labelled nuclei prior to FANS.**

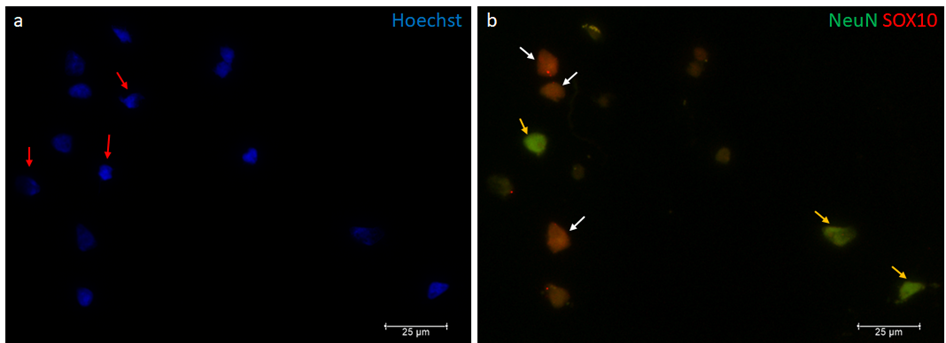

**(a)**

**(b)**

Immunocytochemistry images (40× magnification) show NeuN-labelled nuclei (green), SOX10-labelled nuclei (red), and Hoechst counterstain (blue). Images were acquired using Leica XS software. Scale bar = 25 µm. (**a**) Hoechst staining highlights the total nuclei population. Red arrows indicate nuclei negative for both NeuN and SOX10, which therefore do not appear in the antibody fluorescence channels. (**b**) Merged fluorescence channels reveal a mixed population of NeuN⁺ nuclei (yellow arrows) and SOX10⁺ nuclei (white arrows) in the unsorted sample.

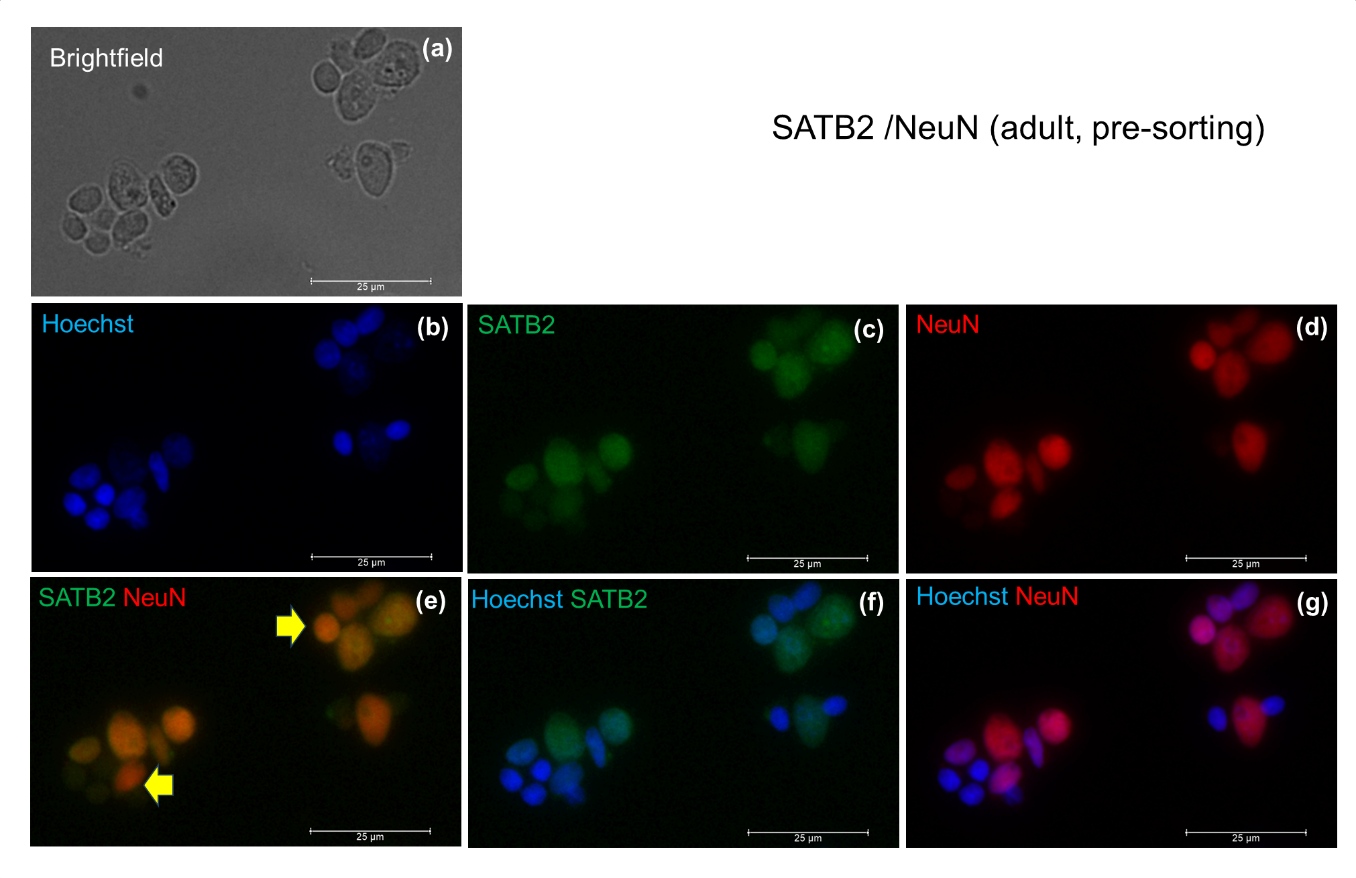

**Figure S5. Validation of neuronal nuclei antibody staining.** Immunocytochemistry images (40× magnification) showing SATB2 (green), NeuN (red), and Hoechst counterstain (blue) in unsorted nuclei. Images were acquired and analysed using Leica XS software. Scale bar = 25 µm. (**a**) Brightfield image showing the localisation of nuclei. (**b**) Hoechst staining of all nuclei. (**c**) SATB2 fluorescence channel. **(d**) NeuN fluorescence channel. **(e**) Overlay of SATB2 and NeuN signals highlights nuclei that are SATB2⁻/NeuN⁺ (yellow arrows). (**f–g**) Comparison of Hoechst/SATB2 (**f**) and Hoechst/NeuN (**g**) overlays confirms that only a subset of NeuN⁺ neurons also express SATB2, consistent with SATB2 marking a specific excitatory neuronal subpopulation.

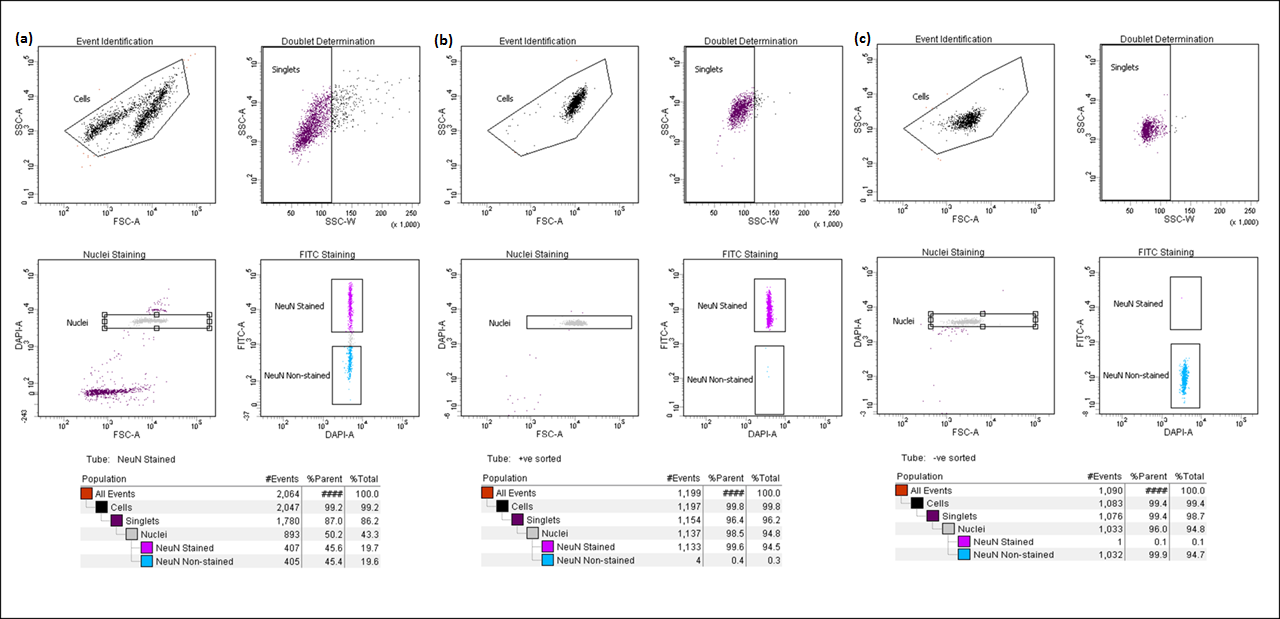
**Figure S6. Validation of sorting purity by re-analysis of NeuN-labelled nuclei.**

To evaluate the accuracy of the FANS gating strategy and confirm the purity of sorted fractions, aliquots of NeuN⁺ (neuron-enriched) and NeuN⁻ (glia-enriched) nuclei were re-analysed immediately after sorting. (**a**) Representative FANS plot from a postnatal human cortical sample showing the initial NeuN–Alexa Fluor 488 gating used to define NeuN⁺ and NeuN⁻ populations. (**b**) Re-analysis of the NeuN⁺ sorted nuclei demonstrated high purity, with nearly all events falling within the NeuN⁺ gate. (**c**) Re-analysis of the NeuN⁻ fraction similarly confirmed minimal contamination by NeuN⁺ events. These results validate the specificity of NeuN staining and demonstrate the reliability of the FANS workflow for generating highly purified neuronal and non-neuronal nuclear populations.

**Figure S7. Gene expression validation confirms cell-type specificity of FANS-sorted nuclei.**

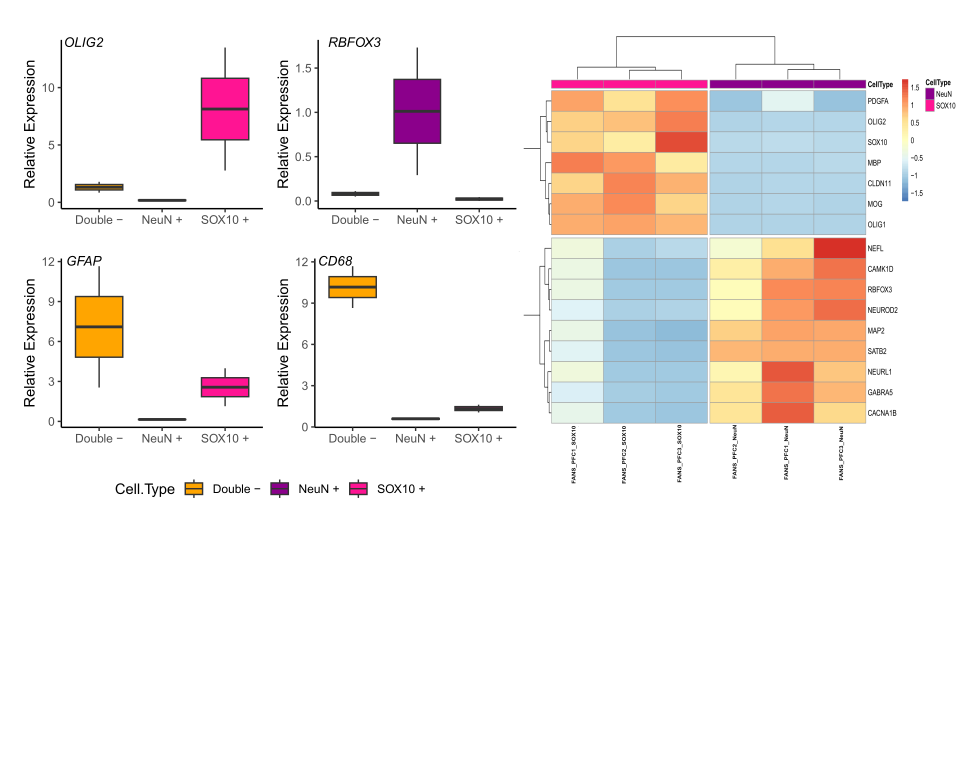

**(a)**

**(b)**

(**a**) Relative qPCR analysis of canonical marker genes in FANS-sorted nuclei. Expression of each marker was quantified relative to the geometric mean of five housekeeping genes (*ACTB*, *EIF4A2*, *GAPDH*, *SF3A1*, and *UBC*) and normalised to the unsorted (bulk) nuclei fraction using the ΔΔCt method. *RBFOX3* (neuronal marker) was highly enriched in the NeuN⁺ population; *OLIG2* (oligodendrocyte marker) showed strong enrichment in the SOX10⁺ fraction; and *GFAP* (astrocyte marker) and *CD68* (microglial marker) were elevated in the NeuN⁻/SOX10⁻ (double-negative) population, consistent with expected cellular identities. (**b**) Bulk RNA-seq performed on NeuN⁺ and SOX10⁺ nuclei (n = 3 per fraction) further confirmed cell-type specificity. A heatmap of selected marker genes demonstrates clear transcriptional separation between neuronal and oligodendrocyte-enriched populations.

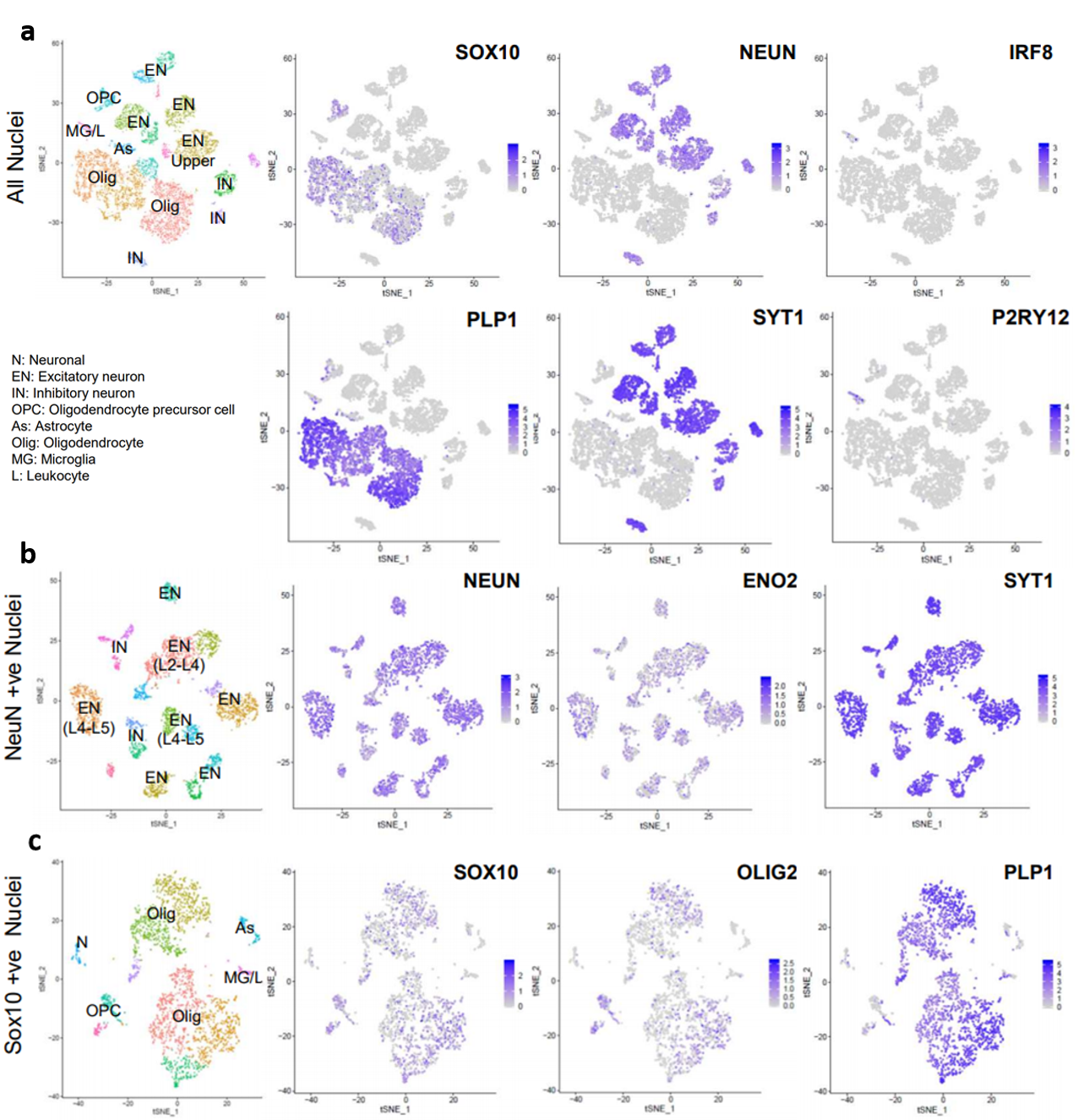

**(a)**

**(b)**

**(c)**

**Figure S8. Single-nucleus RNA-seq of FANS-sorted nuclei from human prefrontal cortex using 10x Genomics.**

(**a–c**) t-SNE visualisations in which each dot represents an individual nucleus profiled using 3′ snRNA-seq. (**a**) Combined dataset of 9,448 nuclei, including total (unsorted) nuclei from two individuals, SOX10⁺ nuclei from two individuals, and NeuN⁺ nuclei from one individual. Expression of canonical marker genes is shown: SOX10 and PLP1 (oligodendrocytes), NEUN and SYT1 (neurons), and IRF8 and P2RY12 (microglia). (**b**) Subset of 1,642 NeuN⁺ nuclei from one donor showing expression of neuronal markers NEUN, ENO2, and SYT1. (**c**) Subset of 2,649 SOX10⁺ nuclei from two donors showing expression of oligodendrocyte lineage markers SOX10, OLIG2, and PLP1.

Manual cell-type annotation identified neuronal (N), excitatory neuron (EN), inhibitory neuron (IN), oligodendrocyte precursor (OPC), astrocyte (As), oligodendrocyte (Olig), and microglial (MG) clusters. The clear concordance between expression of sorting markers (SOX10, NEUN, IRF8) and established cell-type marker genes confirms the specificity and accuracy of the FANS-based nuclei isolation.

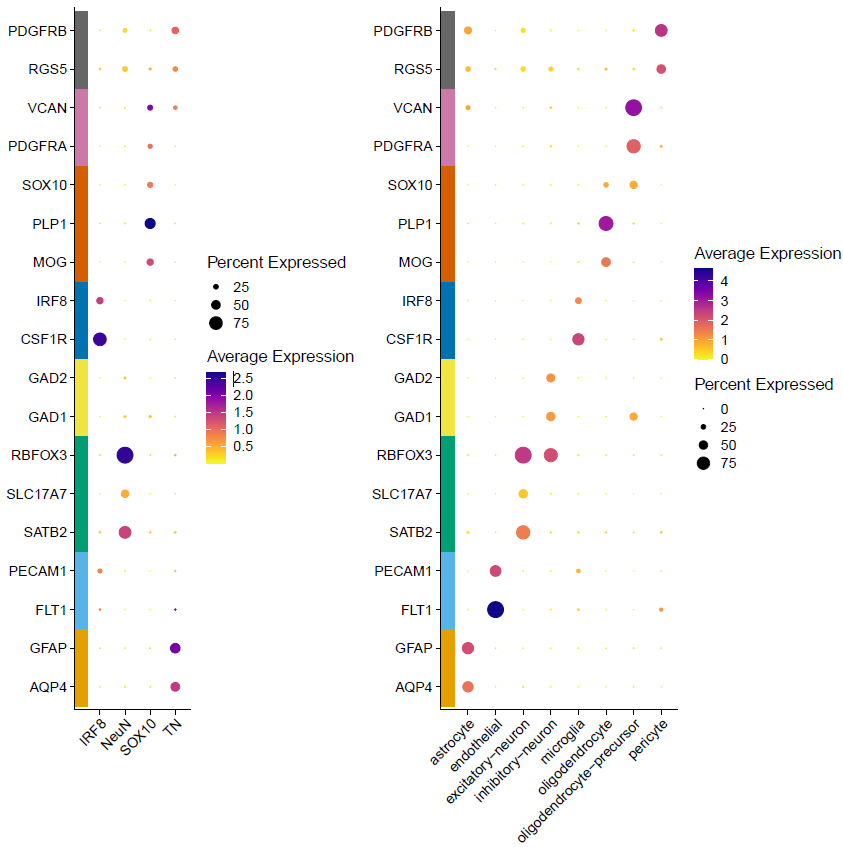

**(b)**

**(a)**

**Figure S9. Parse single-nucleus RNA-seq of FANS-sorted nuclei from human prefrontal cortex.**

Approximately 100,000 nuclei per sample, isolated using our FANS protocol, were processed for single-nucleus RNA sequencing (snRNA-seq) using the Parse Biosciences Evercode™ Low-Input Nuclei Fixation kit. Four FANS-sorted populations were collected from each of four individuals: NeuN⁺ (neuron-enriched), Sox10⁺ (oligodendrocyte-enriched), IRF8⁺ (microglia-enriched), and triple-negative (TN = NeuN⁻/Sox10⁻/IRF8⁻).

(a) Dot plot showing log-normalised mean/scaled expression of key marker genes across the four sorted populations, demonstrating that each group is enriched for its expected cell-type markers. (b) Expression of the same marker genes displayed across cell-type annotations derived from the Seurat analysis pipeline.
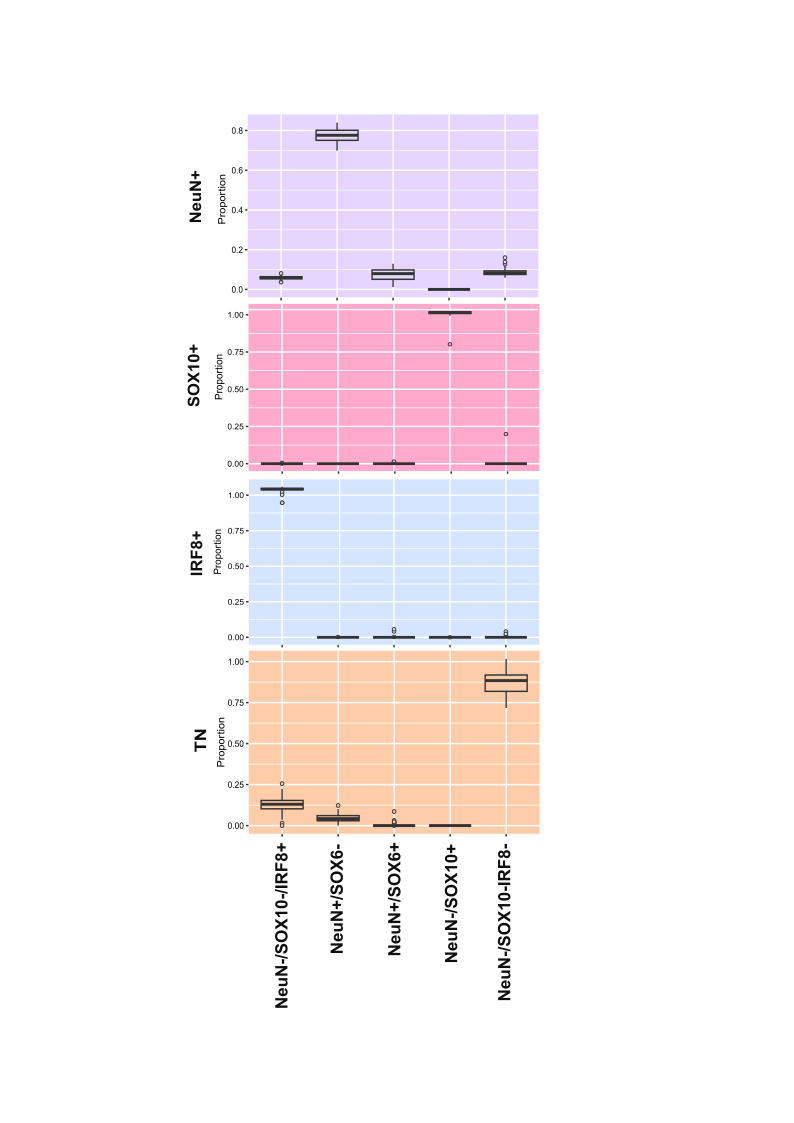
**Figure S10. Estimated cellular proportions for each sample grouped by FANS-sorted fraction.**

Cell-type proportions were estimated using the CETYGO package in R, applying the ANOVA model and the Panel 8 reference dataset, which includes reference profiles for NeuN⁺, SOX10⁺, IRF8⁺, and SOX6⁺ nuclei. SOX6 serves as a marker of inhibitory neurons. The predicted proportions closely match the expected cellular identities for each FANS-sorted fraction, confirming the effectiveness of the sorting strategy

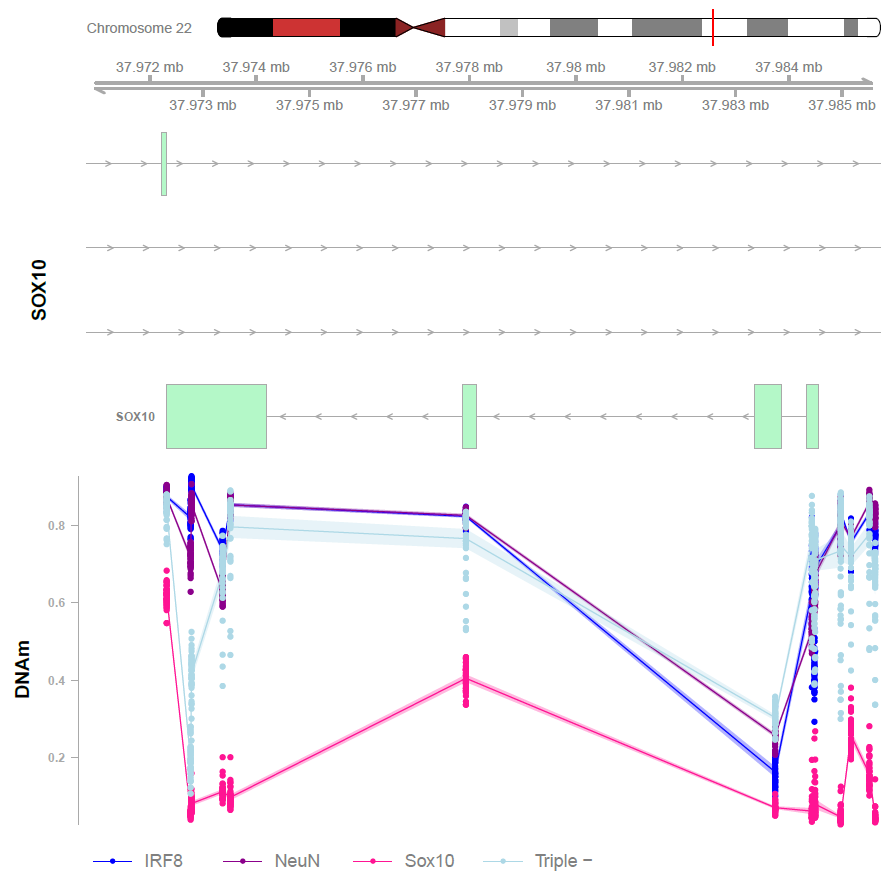

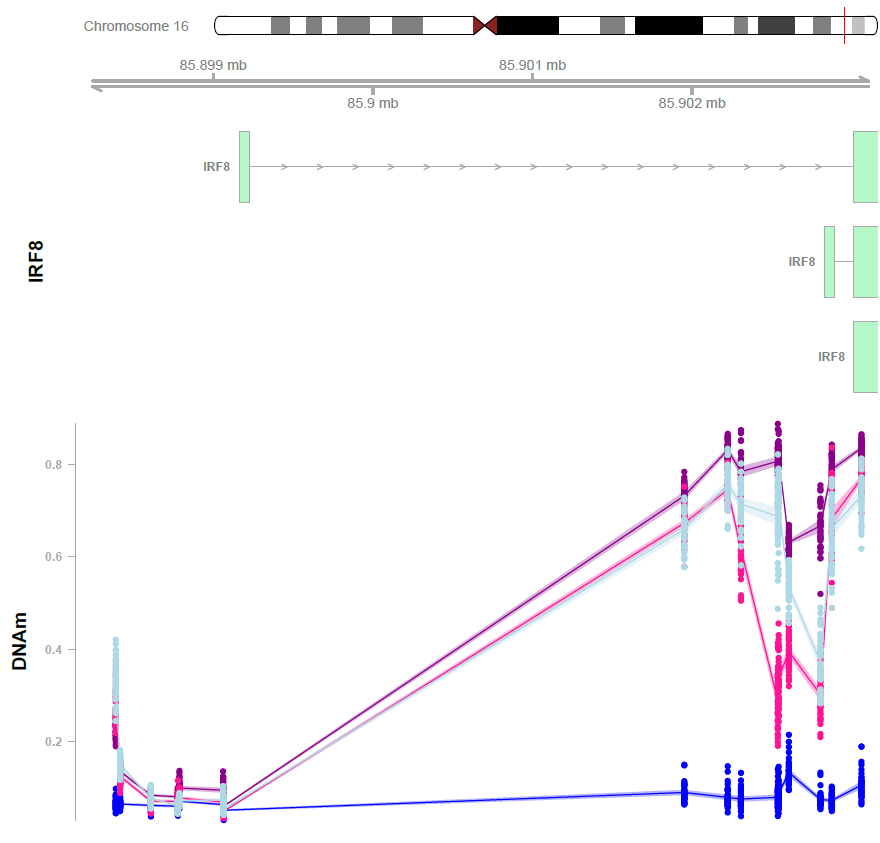

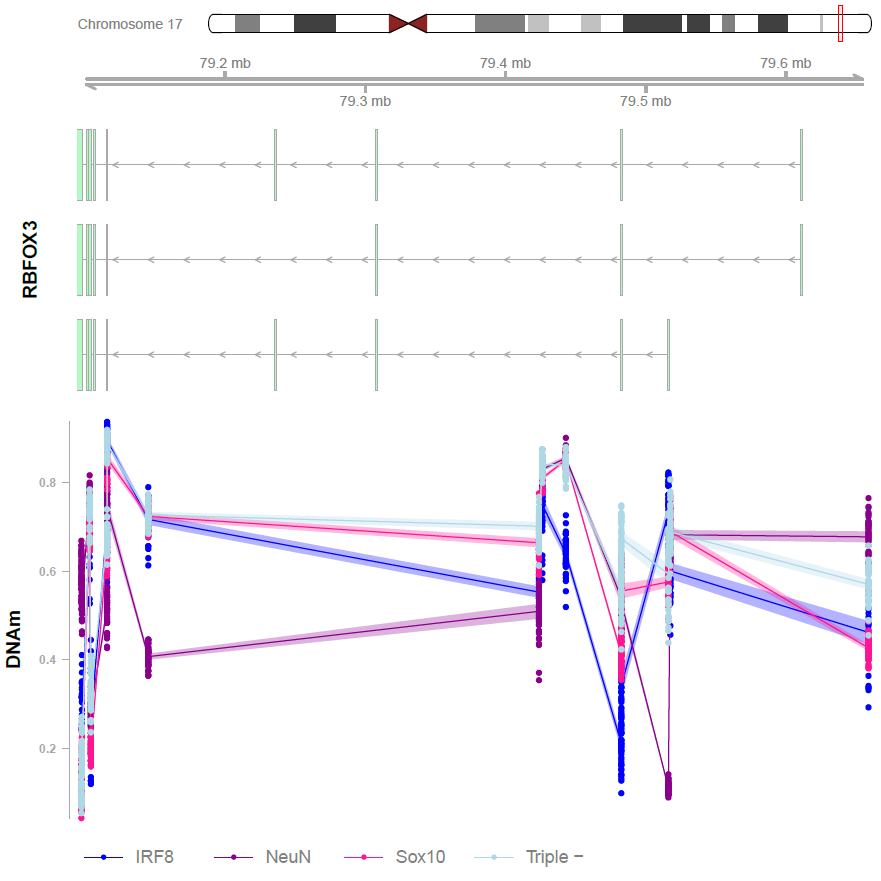

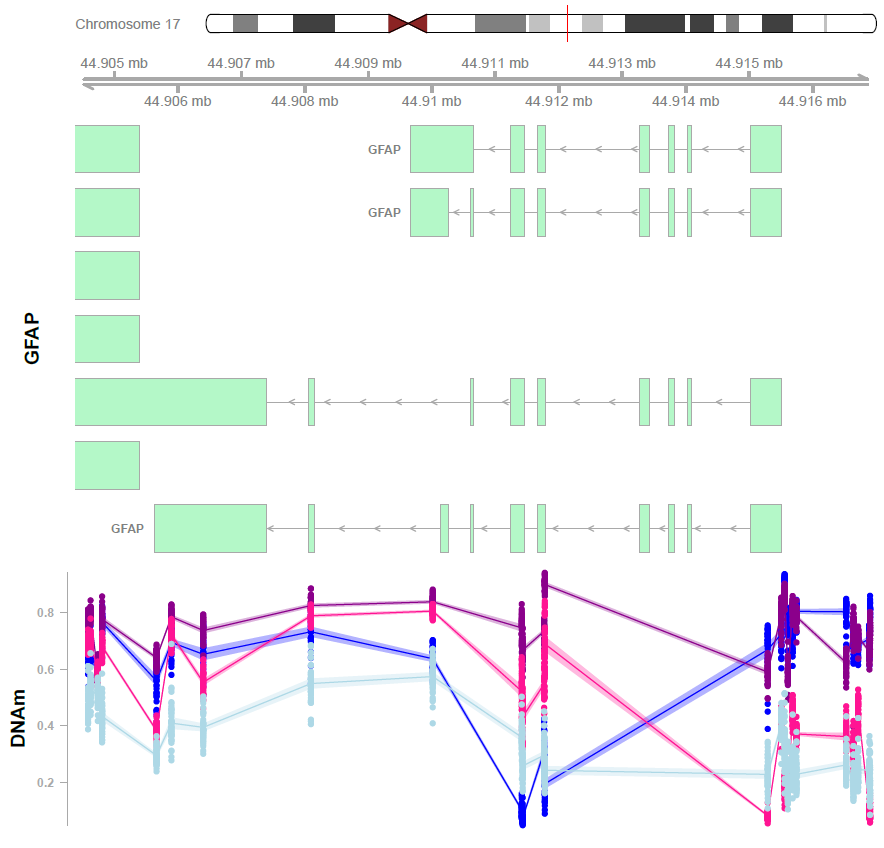

**(a)**

**(b)**

**(c)**

**(d)**

**Figure S11. DNA methylation array profiling reveals cell-type-specific patterns in FANS-sorted nuclei.**

(a–d) Gene-level DNA methylation profiles for representative marker genes demonstrate distinct methylation signatures across the four major FANS-isolated nuclei populations. (a) RBFOX3 (NeuN), a neuronal marker, shows reduced methylation specifically in the NeuN⁺ population relative to the other fractions. (b) SOX10⁺ nuclei (oligodendrocyte-enriched) exhibit characteristic hypomethylation across the SOX10 locus compared with the remaining populations. (c) IRF8⁺ nuclei (microglia-enriched) show pronounced hypomethylation at CpG sites across the IRF8 gene. (d) The triple-negative population (NeuN⁻/SOX10⁻/IRF8⁻) displays hypomethylation across GFAP, consistent with enrichment for astrocyte-lineage nuclei.

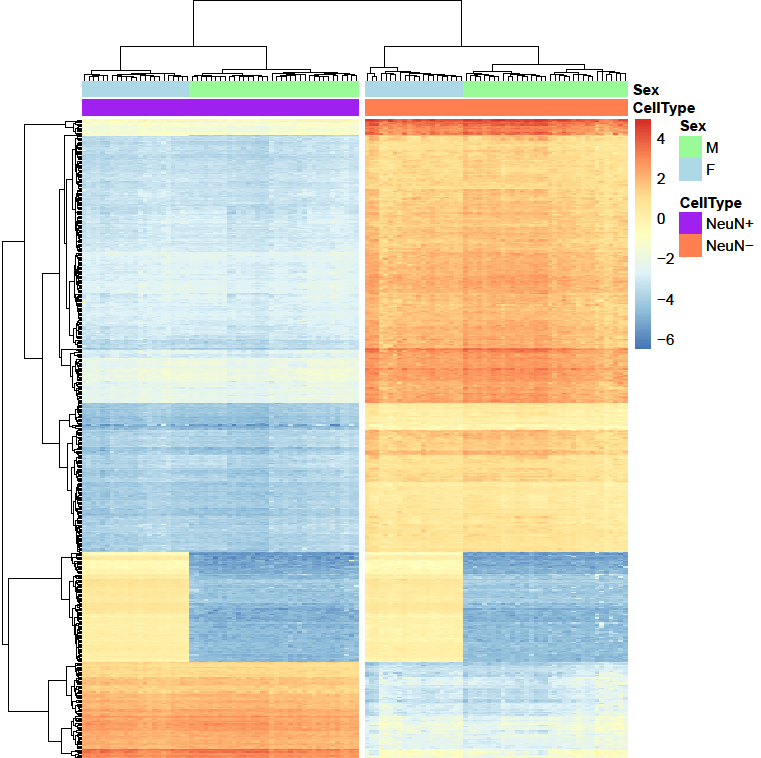

**Figure S12. DNA methylation differences between neuronal and non-neuronal nuclei in mouse cortex.**

Heatmap of the top 1,000 most variable differentially methylated positions (DMPs) across FANS-sorted nuclei from mouse cortex. NeuN⁺ nuclei (purple), representing neuron-enriched populations, exhibit distinct DNA methylation profiles compared with NeuN⁻ nuclei (orange), which represent non-neuronal populations.

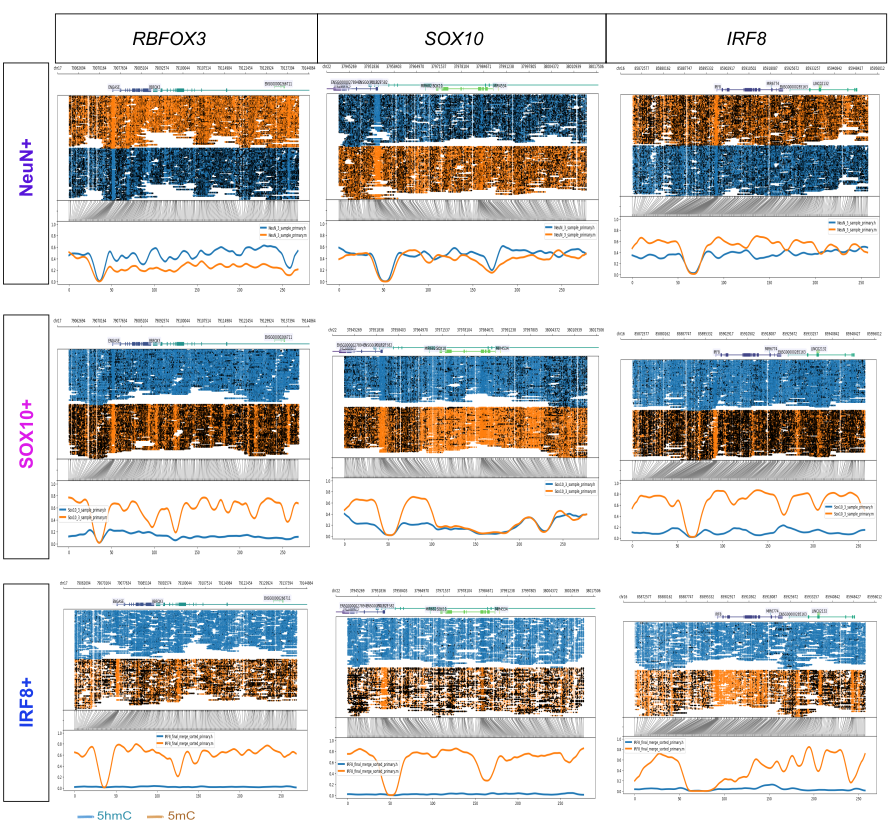

**Figure S13. Oxford Nanopore sequencing reveals cell-type-specific DNA modification profiles.**

Genome-wide Oxford Nanopore Technologies (ONT) sequencing of DNA from three FANS-sorted nuclei populations demonstrates distinct patterns of 5-methylcytosine (5mC) (orange) and 5-hydroxymethylcytosine (5hmC) (blue) across representative cell-type marker genes. These modification differences highlight the unique epigenetic landscapes of each neuronal, oligodendrocyte, and microglial population.

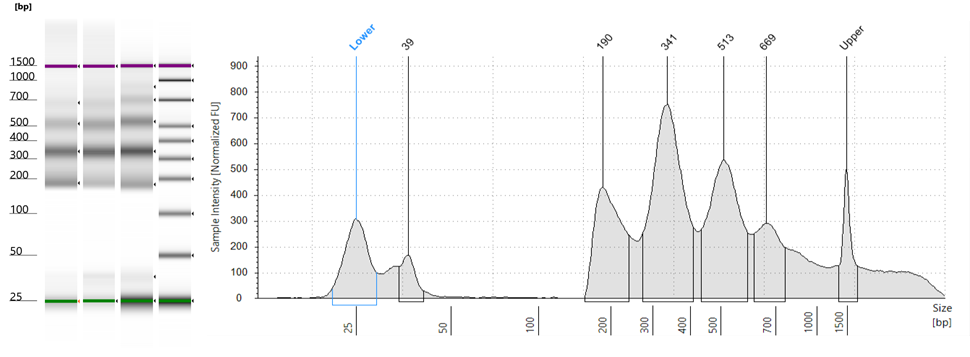

**Figure S14 Representative trace of ATAC-seq libraries**

Visualised using an Agilent D1000 HS ScreenTape. The peaks of the trace are indicative of the periodicity of the chromatin structure and show nucleosome-free, mononucleosome, dinucleosome and multinucleated fragments. The peaks occur at a frequency of 150-180bp, concordant with the length of DNA wrapped around each nucleosome.

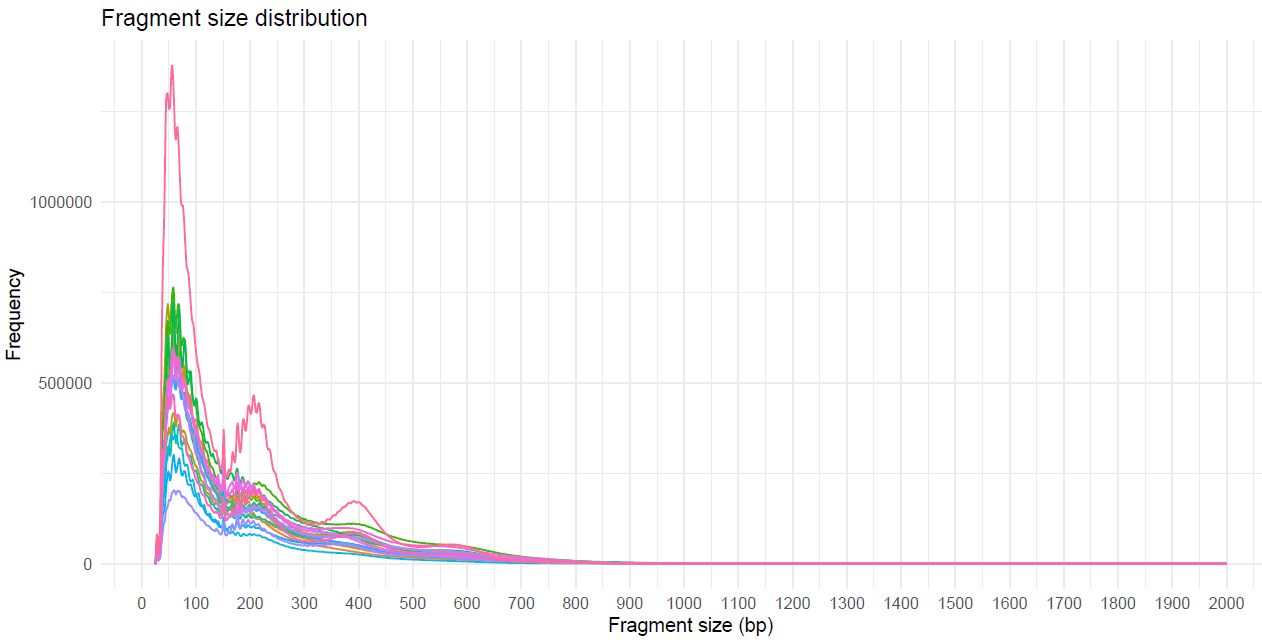

**Figure S15. ATAC-seq fragment length profiles from FANS-isolated nuclei.**

ATAC-seq libraries generated from FANS-sorted nuclei exhibit the expected fragment length distribution, with clear enrichment at ~100 bp (nucleosome-free fragments) and at ~200 bp and ~400 bp (mono- and di-nucleosome–associated fragments). This characteristic pattern confirms high-quality chromatin accessibility profiling from the sorted nuclei.

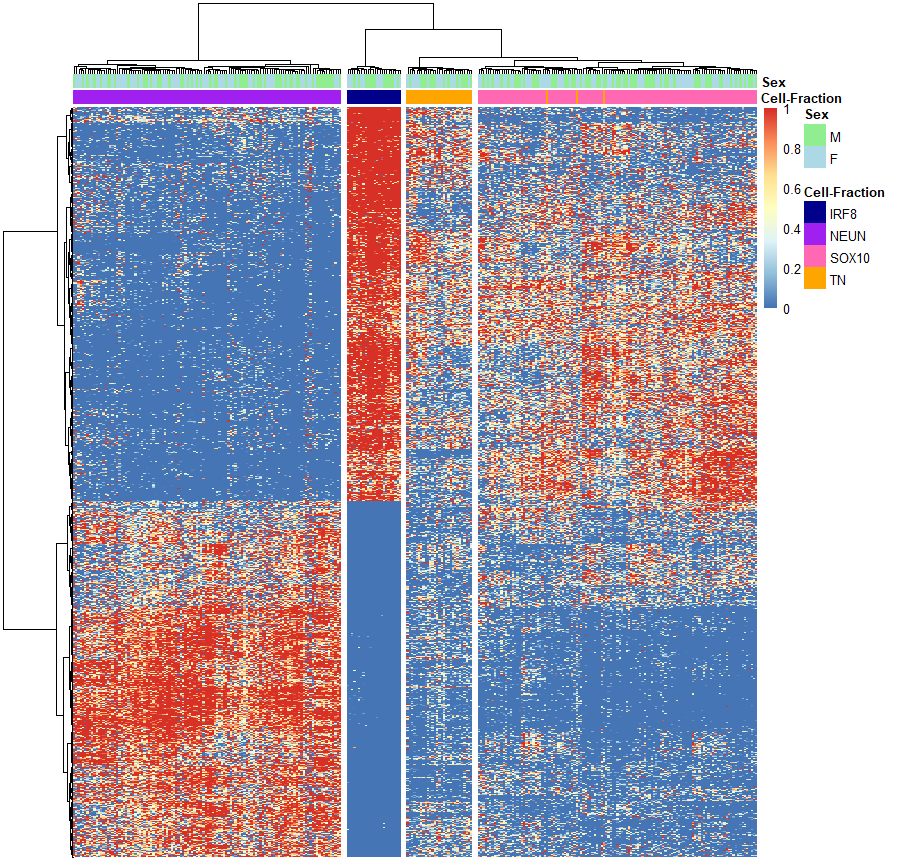

**Figure S16**. **ATAC-seq data from FANS-isolated nuclei**.

Hierarchical clustering of chromatin accessibility profiles across samples, based on the top 5,000 most variable ATAC-seq peaks across all cell types, demonstrates clear separation of the four major FANS-isolated nuclear populations according to their genome-wide accessibility signatures. Each row represents a single ATAC-seq peak, and each column corresponds to an individual sample. Chromatin accessibility values were normalised and scaled; blue indicates lower accessibility and red indicates higher accessibility.

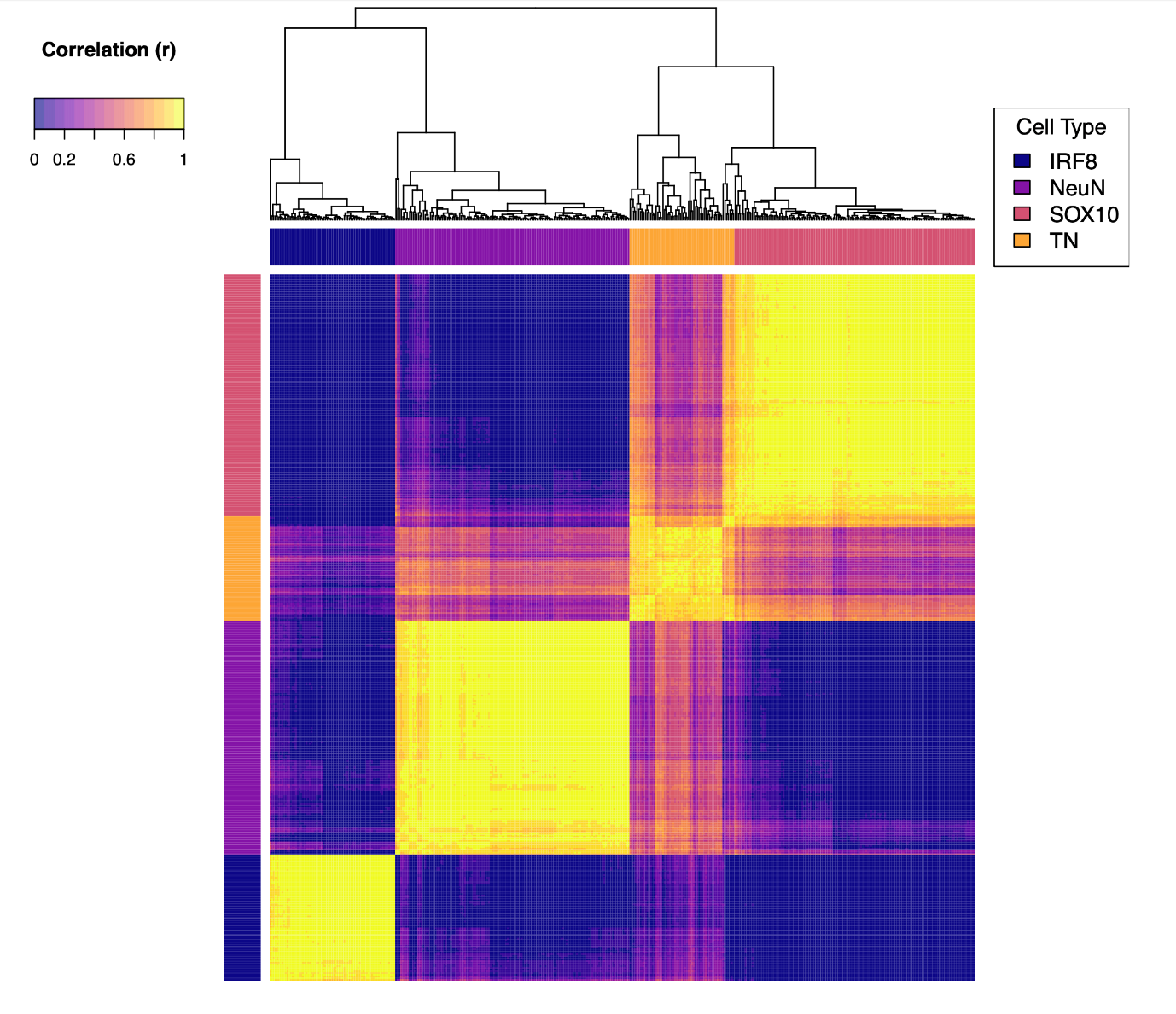

**Figure S17. H3K27ac CUT&Tag profiling of FANS-sorted nuclei.** (A) Hierarchical clustering of Pearson correlation coefficients (r) across samples, calculated using variance stabilising transformed counts in the top 5,000 most variable H3K27ac peaks combined across all cell types. Samples cluster according to their FANS-sorted population, demonstrating strong cell-type-specific enhancer signatures.
