## Supplementary Table 1 for "Optimised fluorescence-activated nuclei sorting for epigenomic analysis of cortical cell types"

| *Supplementary Table S1.* Pre-optimised TaqMan assays for targeted gene expression analysis (Life Technologies). | | | | |
| --- | --- | --- | --- | --- |
|  | **Gene Symbol** | **Gene Name** | **TaqMan Assay ID** | **Category** |
| 1 | *ACTB* | actin beta | Hs99999903_m1 | housekeeping |
| 2 | *EIF4A2* | eukaryotic translation initiation factor 4A2 | Hs00756996_g1 | housekeeping |
| 3 | *GAPDH* | glyceraldehyde-3-phosphate dehydrogenase | Hs99999905_m1 | housekeeping |
| 4 | *SF3A1* | splicing factor 3a subunit 1 | Hs01066327_m1 | housekeeping |
| 5 | *UBC* | ubiquitin C | Hs00824723_m1 | housekeeping |
| 6 | *RBFOX3* | RNA binding protein, fox-1 homolog 3 | Hs01370654_m1 | target gene |
| 7 | *ENO2* | enolase 2 | Hs00157360_m1 | target gene |
| 8 | *GFAP* | glial fibrillary acidic protein | Hs00909233_m1 | target gene |
| 9 | *OLIG2* | oligodendrocyte lineage transcription factor 2 | Hs00377820_m1 | target gene |
| 10 | *CD68* | CD68 molecule | Hs00154355_m1 | target gene |
